# Enhancing Uterine Receptivity for Embryo Implantation through Controlled Collagenase Intervention

**DOI:** 10.1101/2022.01.18.476712

**Authors:** Eldar Zehorai, Tamar Gross, Elee Shimshoni, Ron Hadas, Idan Adir, Ofra Golani, Guillaume Molodij, Ram Eitan, Karl E. Kadler, Orit Kollet, Michal Neeman, Nava Dekel, Inna Solomonov, Irit Sagi

**Author notes:** Equal contribution (Eldar Zehorai, Tamar Gross). Corresponding authors: Irit Sagi, PhD, Professor, Department of Biological Regulation, Weizmann Institute of Science, Rehovot 7610001, Israel, Inna Solomonov, PhD, Department of Biological Regulation, Weizmann Institute of Science, Rehovot 7610001, Israel.

## Abstract

Successful embryo implantation within the uterine wall requires intricate endometrial remodeling. Impaired endometrial receptivity, a common cause of infertility, often results from ineffective remodeling processes. Here, we demonstrate that a single dose of human collagenase-1 administered into the mouse uterus enhances embryo implantation rates. Mechanistically, collagenase-1 induces remodeling of the endometrial extracellular matrix (ECM), leading to the degradation of collagen fibers and proteoglycans. This process releases matrix-bound bioactive factors, such as VEGF, which facilitates local vascular permeability and angiogenesis. Furthermore, collagenase-1 treatment increases NK cell infiltration and elevates levels of the cytokine LIF, a key factor in embryo implantation. Remarkably, the overall structural integrity of the uterine tissue remains uncompromised, even in the presence of reduced tension in endometrial collagen fibers. To assess pre-clinical potential, in-uteri application of collagenase-1 successfully rescued implantation in mouse models subjected to heat stress and embryo transfer, conditions known for their adverse impact on implantation rates. Importantly, ex-vivo exposure of human uterine tissue to collagenase-1 induced collagen de-tensioning and the release of VEGF, demonstrating similar processes observed in the mouse settings, and the potential relevance of this treatment to human conditions. Our findings underscore the immense clinical potential and feasibility of collagenase treatment to enhance uterine receptivity for embryo implantation, offering a controlled and minimally invasive intervention. This innovative approach not only demonstrates the potential to enhance efficiency in livestock breeding but, more importantly, signifies a substantial promise in supporting medical interventions for human reproduction in clinical settings.

## Introduction

Embryo implantation is a complex process involving dynamic genetic, biophysical, and biochemical rearrangements in the uterine endometrium layer. A prerequisite for successful implantation is the establishment of an appropriate endometrial microenvironment, influenced by many parameters such as hormone production, extracellular matrix (ECM) remodeling, local immune-system adjustment, and angiogenesis^1-3^. Proper ECM remodeling is particularly crucial for creating optimal receptive conditions and synchronization of functionally appropriate embryo-endometrium crosstalk. This process takes place during each reproductive cycle to prime the endometrium for implantation and continues during the window of implantation (WOI) and the decidualization phase^4^.

During embryo implantation, the spatiotemporal expression of abundant ECM components in the endometrium is altered^5-7^. In mice, collagen types I and III are locally reduced at the implantation sites due to extensive remodeling^6,7^. Similarly, the ultra-structure of fibrillar collagens in the human uterus changes during the first trimester of pregnancy. These observations suggest that fibrillar collagens play a role in trophoblast cell adhesion and invasion^8^, and impaired remodeling of endometrial extracellular proteins correlates with implantation failure and pregnancy loss.

Endometrial remodeling during embryo implantation is mediated by various enzymes, particularly matrix metalloproteinases (MMPs)^9^, which are widely expressed by both the endometrium and trophoblast cells^10,11^. MMPs regulate trophoblast invasion and contribute to endometrial vascular remodeling and inflammatory processes by proteolysis of ECM molecules and cell receptors^12-14^. Exposing trophoblasts ex-vivo to gelatinases (MMP-2/9) did not increase the pregnancy rate but slightly improved pups’ viability, implying an unelucidated mechanism involving collagen reduction in the blastocysts^15^. However, the potential of MMP-mediated collagen/ECM remodeling mechanisms in promoting normal uterine tissue physiology, such as implantation, has been largely unexplored. Addressing the challenge of improving conditions for successful implantation through optimized endometrial ECM receptive state may be facilitated by the precise regulation and understanding of endometrial remodeling. This approach preserves trophoblasts from ex-vivo interventions.

Previously, we reported that collagenase-1 (MMP-1)-mediated proteolysis of collagen-rich ECM leads to unique changes in collagen morphology, ECM viscoelastic properties, molecular composition and cell adhesion capacity. These alterations in the ECM have profound impacts on neighboring cells, affecting their morphology, signaling patterns and gene-expression profiles. Notably, we observed enhanced adhesion and invasion of fibroblasts in collagenase-1-treated collagen-rich ECM in an *ex vivo* settings^16^. These findings motivated us to investigate the effects of mild ECM remodeling facilitated by collagenase-1 treatment on embryo implantation rates.

Here, we demonstrate that a single topical in-uterus administration of collagenase-1 significantly improves embryo implantation rates in mice, regardless of genetic background. Collagenase-1 treatment induces mild extracellular remodeling by reducing the tension (de-tensioning) of endometrial collagen fibers, releasing matrix-bound active factors, promoting neo-angiogenesis, and initiating a pro-inflammatory environment, all necessary for successful embryo implantation. Our study highlights the potential utilization of in-uterus collagenase-1-specific proteolysis for increasing embryo implantation rates and provide new mechanistic insights into endometrial receptivity.

## Results

### Endometrial collagen remodeling boosts embryo implantation

We previously reported that collagenase-1-induced proteolysis of natural collagen fascicles from rat tendons results in specific and unique ECM changes^16^. Further examination of the collagen network by scanning electron microscopy (SEM), revealed less densely packed, relaxed and disorganized arrangement of collagen fibrils within each fiber. These observations signify that collagen fibers undergo de-tension when treated with collagenase-1 (Fig. 1A). In addition, we showed that fibroblasts exhibit enhanced adhesion to collagen-rich ECM that has been pretreated with collagenases compared to untreated collagen^16^. Based on these results and recognizing the crucial role of endometrial ECM-remodeling by MMPs for successful embryo implantation^9^, we hypothesized that inducing moderate remodeling of collagen fibers through exogenous collagenase-1 treatment would enhance embryo-endometrium adherence and invasion, thereby improving the implantation process. To test this hypothesis, we incubated five mouse blastocysts *ex vivo* with native collagen-rich ECM pretreated with either vehicle or collagenase-1. After 4 hours of incubation, most of the blastocysts (4 out of 5) interacted with the collagenase-1-treated collagen-rich ECM while none interacted with the vehicle-treated matrix (Fig. 1B). This observation that collagen/ECM proteolysis enhances blastocyst adherence and interactions *ex vivo*, prompted us to investigate the effect of in-uterus collagenase-1 treatment on embryo implantation *in vivo*. For this purpose, we utilized a spontaneous pregnancy model in which copulated ICR female mice were topically administered with a low dose of recombinant collagenase-1 (1.25 µg) or vehicle at day E2.5 (2.5 days post-coitum) (Fig. 1C). The number of embryo implantation sites, recorded in the harvested uteri 4 days later at E6.5, showed a significant increase of ∼60% in collagenase-1-treated mice (Fig. 1D). This effect was also observed in C57BL/6 mated females, demonstrating a significant improvement of ∼50% in embryo implantation (Fig. S1). Notably, the litter of treated animals was healthy and viable, exhibiting normal development and behavior. Additionally, the pups weight measured 3 weeks after birth was within the expected range of average weight reported for healthy mice of the same age and strain (12±3 g)^17^.

**Fig. 1.**
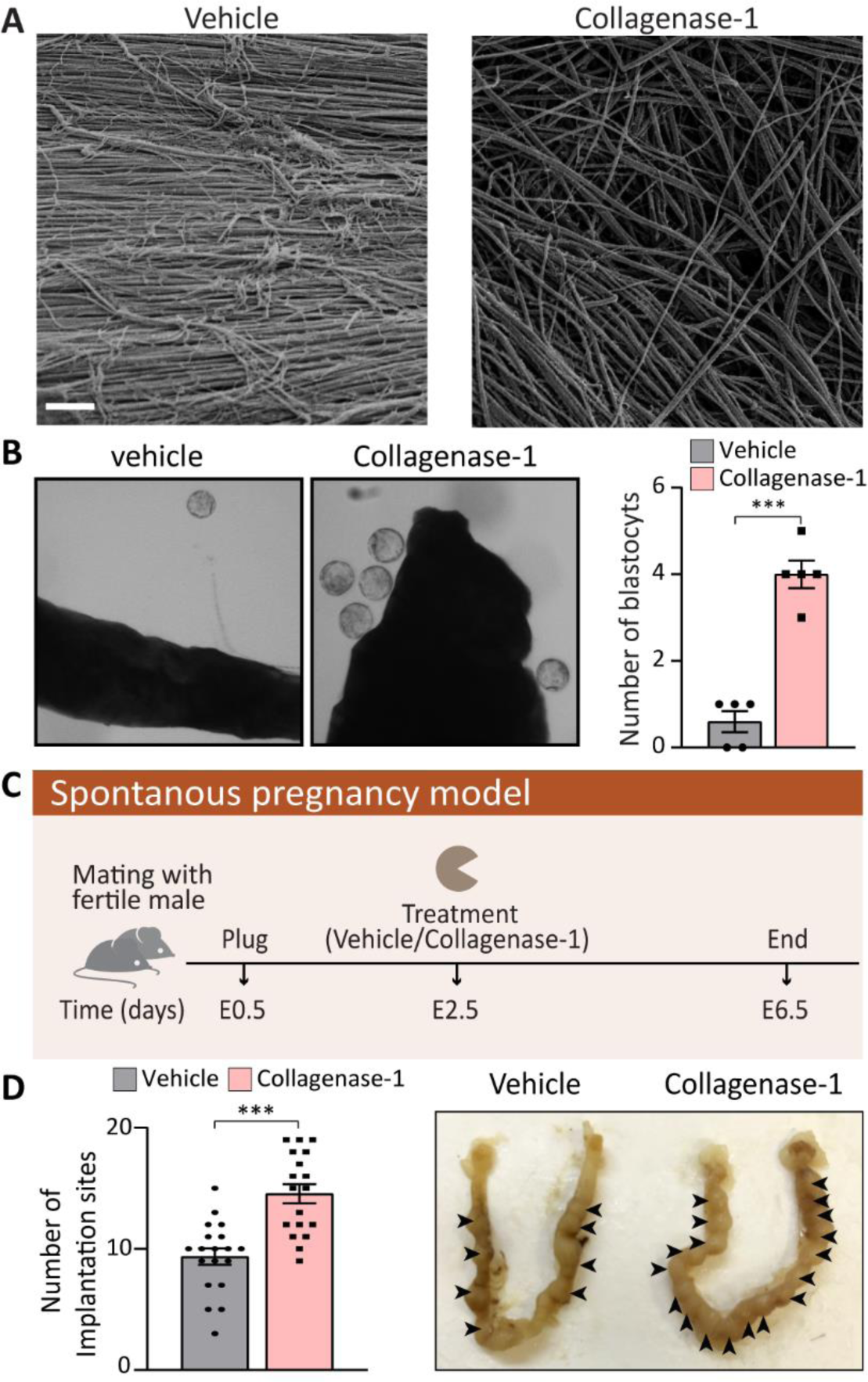
Collagenase-1 improves the adherence of embryos ex vivo and implantation rate in vivo. **(A)**. Representative SEM images of native rat tail collagen fiber pretreated with either collagenase-1 or vehicle, demonstrating alterations in fibrils alignment within the fiber due to collagenase-1 treatment (scale-2µm). (**B**) Representative images of blastocyst adherence to fascicle-derived native collagen-rich ECM pretreated with collagenase-1 or vehicle. Blastocysts that were in close proximity to the collagen were counted (n = 5). (**C**) Schematic representation of the spontaneous pregnancy model. (**D**) Quantification of implantation sites and representative images of the uteri at E6.5. Implantation sites marked by arrowheads (n = 30). Data were analyzed by an unpaired, two-tailed t-test. Results are presented as mean ± SEM with significance: ***p < 0.001.

### Collagenase-1 treatment maintains uterine tissue integrity

To elucidate the molecular mechanisms underlying collagenase-1’s enhancement of embryo-ECM interactions, we examined the overall structure of the endometrial tissue at E4.5, a stage at which the embryo is fully attached and decidualization has commenced. No discernible differences were observed in the uterus tissue layers comparing collagenase-1 treated and control groups. The uteri myometrium and endometrium layers appeared intact in both groups with clearly demarcated borders (Fig. 2A). Pathological examination further confirmed that collagenase-1 treatment did not induce any cellular abnormalities or distinct morphological changes in epithelia, stroma, and myometrial smooth muscle cells of the uterus. To exclude the possibility that collagenase-1 treatment induces pathological trophoblast invasion or excessive embryo invasiveness into the myometrium, we imaged the implantation site of GFP-expressing embryos within the uterine fibrillar collagens using two-photon microscopy with a second-harmonic generation (SHG) modality. This examination was performed at E6.5 when the invasion process is completed, and gastrulation begins. Representative images in Figure 2B depict GFP-expressing embryos interacting solely with endometrial collagens. Thus, using a low dose of collagenase-1 did not lead to abnormal invasion depth of the implanted embryos nor did it interfere with overall uterine tissue integrity.

**Fig. 2.**
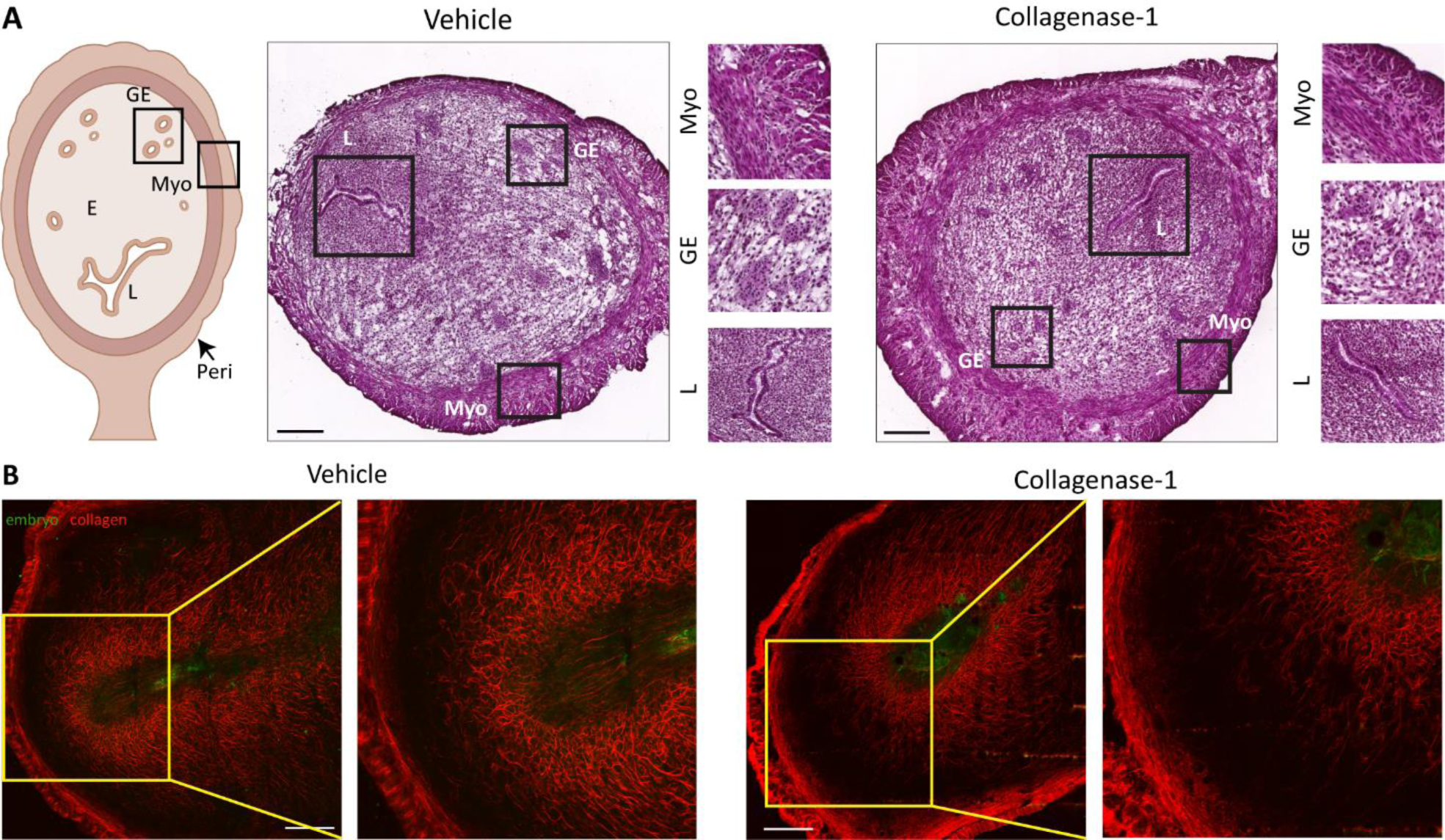
Topical administration of collagenase-1 preserves tissue integrity, avoiding excessive invasiveness of the embryo. **(A)** Schematic and representative H&E images of uteri at E4.5 (n = 3). The endometrium (E), myometrium (Myo), and perimetrium (Peri) layers of the uteri appear intact in both groups. The treatment did not affect the luminal epithelium (L) morphology, which is organized as a simple-columnar monolayer with basal nuclei. The glandular epithelium (GE) remained intact in the treated uteri, and appeared similar to the vehicle-treated uteri. Enlarged insets are shown in the right panels. (**B**) Representative SHG images of transverse uteri sections of vehicle and collagenase-1 treated mice at E6.5, show that collagenase-1 treatment did not affect embryo invasiveness. Enlarged insets are shown at the right panels. green: venus-embryo, red: collagen fibers, scale bar = 50 µm, n = 1.

### De-tensioning of endometrial collagen fibers

Next we profiled collagen organization before and during the window of implantation (WOI), i.e., E2.5 and E4.5, respectively. First, we characterized the effect of collagenase-1 treatment on the organization of endometrial collagen fibers applying SHG modality, a fast and convenient optical method that allows visualization of collagen fiber assembly within the tissue. At E2.5 (1 hr after the treatment) and at E4.5, the treated uteri presented loosely packed and wavy fibers. In contrast, the vehicle-treated uteri maintained dense morphology with well-aligned and tightly packed collagen fibers (Fig. 3A, B). To assess the degree of fiber misalignment, we applied an unbiased orientation analysis developed in-house and calculated the probability of collagen fibers orientation expressed in entropy terms (see Methods). This analysis revealed a slight, but significant increase in orientation entropy in treated compared to control uteri, both at 1h after collagenase treatment (E2.5) and E4.5 (Fig. 3C, top), demonstrating de-tensioning of collagen fibers. Nevertheless, analysis of the endometrial area covered by collagen showed a slight, but not significant reduction between the samples, confirming the absence of substantial fiber degradation (Fig. 3C, bottom). Finally, these analyses revealed that collagenase-1 treatment did not affect the structural organization of the myometrial collagen fibers (Fig. S2). High resolution scanning electron microscope (SEM) images of E2.5 uteri taken 1h after collagenase treatment revealed distinct topological features of relaxed endometrium collagen fibers (Fig. 3D). Their surface was covered by non-aligned wavy fibrils, indicating mild collagenolysis, in contrast to the well-aligned and densely packed fibrils in the control group.

**Fig. 3.**
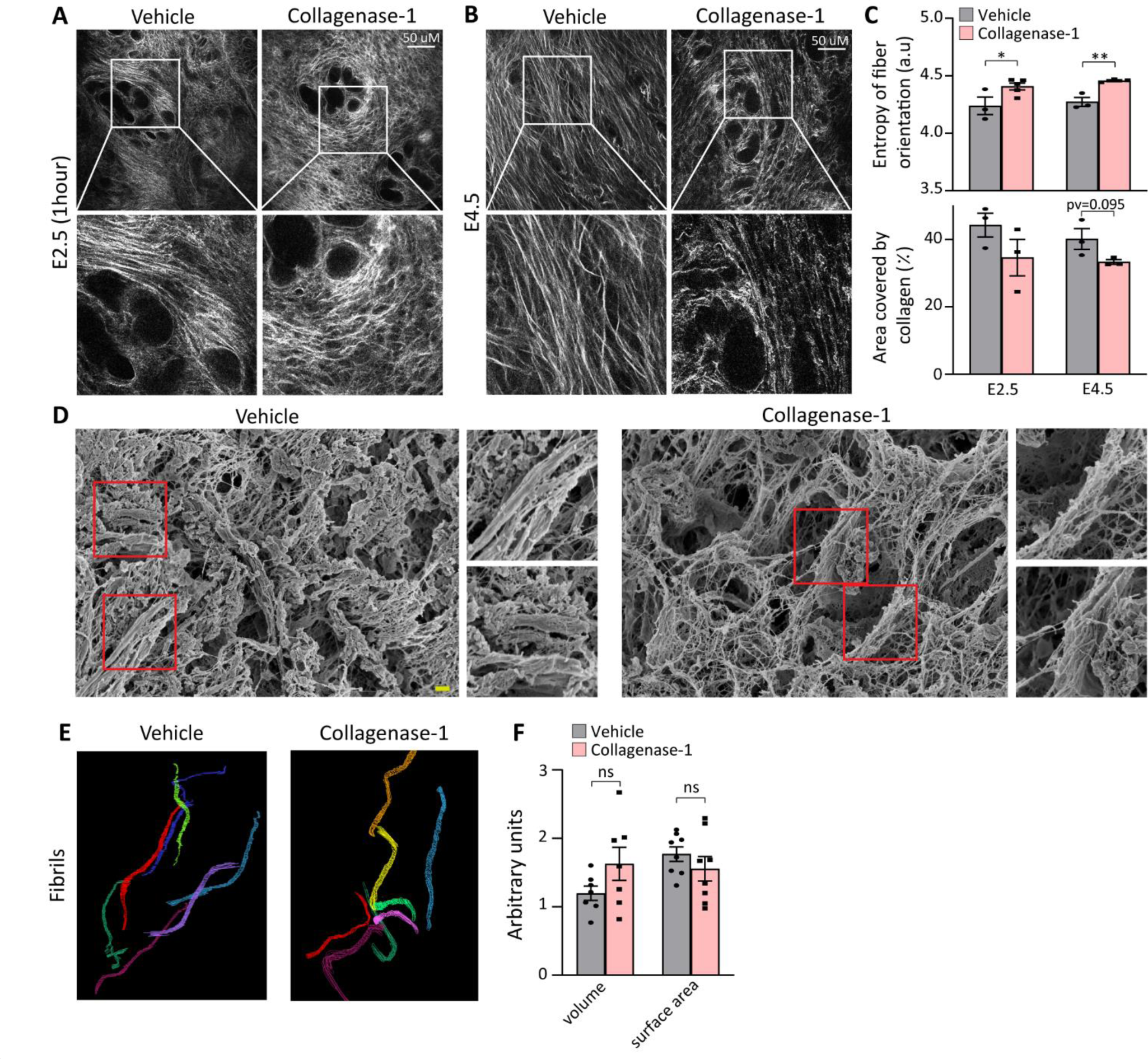
Topical collagenase-1 administration modifies spatial organization of uterine collagen fibrils. **(A, B)** Representative SHG images of longitudinal sections vehicle- and collagenase-1 treated uterine samples at E2.5 (1h after treatment) and E4.5, respectively. Only the endometrium layer is imaged, scale bar = 50 µM, (n≥3). Enlarged insets are shown at the lower panels. **(C)** Measurements of the entropy of fibers orientation (top), and area covered by collagen (bottom). Analysis was done using ImageJ software. **(D)** Representative SEM images of endometrial layers vehicle- and collagenase-1-treated, 1h after treatment. The images display well-aligned dense fibril packing on fiber surfaces of vehicle samples *vs* misaligned wavy fibrils on the fiber surfaces of treated samples, scale bar=1µm. Enlarged insets are shown at the right panels. **(E)** Representative 3D reconstructions from serial block face-Volume EM analysis of endometrium fibrils of vehicle- and collagenase-1-treated uteri. **(F)** Analyses of volume and surface area of fibrils from vehicle- and collagenase-1-treated uteri (n = 8 fibrils). Data were analyzed by an unpaired, two-tailed t-test. Results are presented as mean ± SEM with significance: *p < 0.05, ** p <0.01.

Next, we analyzed the three-dimensional microstructure of endometrium collagen fibrils using a high-resolution serial block-face SEM (SBF-SEM). This volume electron microscopy modality provides sufficient resolution to study individual collagen fibrils by generating a representative reconstructed 3D model^18^. Analysis of randomly selected reconstructed collagen fibrils from samples taken 1 h after treatment did not show significant changes in fibril volume fraction when compared to vehicle samples (Fig. 3E-F), further indicating that collagenase-1 did not induce gross fibril degradation. Taken together, structural analyses at different resolutions provide evidence that mild proteolysis by collagenase-1 treatment mostly manifests in de-tensioning of collagen fibers affecting their spatial organization, rather than intensive fibril degradation. Such collagen proteolysis supports trophoblast invasion and angiogenesis in vivo^8,19^.

### COL1A1 and DCN: collagenase-1 substrates in pregnant uteri

To gain a deeper understanding of the molecular effects of collagenase-1 treatment, we investigated the substrates released by collagenase-1 from the pregnant uterus. E2.5 uteri were subjected to a 4h incubation with collagenase-1, and the resulting supernatants were subjected to LC-MS/MS proteomics analysis (Fig. 4A). The analysis revealed 41 substrates that pass the threshold of: log2(collagenases-1/vehicle)>0.4 and q<0.1 (Fig 4B). Among these substrates, three proteins were identified as matrisome proteins: collagen alpha-1 (Col1a1), decorin (DCN) and annexin A6 (Anxa6) (Fig 4C). To enrich for collagenase-1 matrisome substrates, we replicated the experiment using decellularized E2.5 uteri, which were incubated with collagenase-1 for 24h (Fig. 4D). In this decellularized setup we identified 38 substrates of collagenase-1 that pass the same threshold of: log2(collagenases-1/vehicle)>0.4 and q<0.1 (Fig. 4E). The most significantly upregulated substrates (log2(collagenases-1/vehicle)>2.5) included ECM-structural proteins and proteoglycans such as dermatopontin, lumican, decorin and collagen alpha-1,2 (Fig. 4F). We then conducted a more in-depth analysis of the decorin protein, which was identified as a collagenase-1 substrate in both experimental setups in pregnant uteri. Our analysis revealed that out of 32 peptides that were cleaved, 9 were semi/non tryptic peptides, and among them 3 exhibited favorable cleavage motifs of collagenase-1, specifically leucine and isoleucine, as indicated by MEROPS (https://www.ebi.ac.uk/merops/cgi-bin/pepsum?id=M10.001) (Fig. 4G). Based on these findings, it is likely that collagenase-1 breaks down decorin proteoglycan in the matrix of the pregnant uterus. Decorin is a crucial stabilizer of the collagen fibrillar network, and its proteolysis can contribute to fiber de-tensioning by inducing collagen misalignment. In addition, decorin binds a variety of signaling molecules, and its degradation can modulate the bio-activity of many important factors, including VEGF and others^20^. To investigate it further, we analyzed the supernatant from the decellularized uteri in a western blot with VEGF-A antibody revealing that VEGF-A was released following incubation with collagenase-1 (Fig 4H, S3). These experiments indicate that collagenase-1 not only remodels collagens but also plays a role in breaking down proteoglycans, like decorin, thereby releasing factors from the matrix and making them bioavailable in the tissue.

**Fig. 4.**
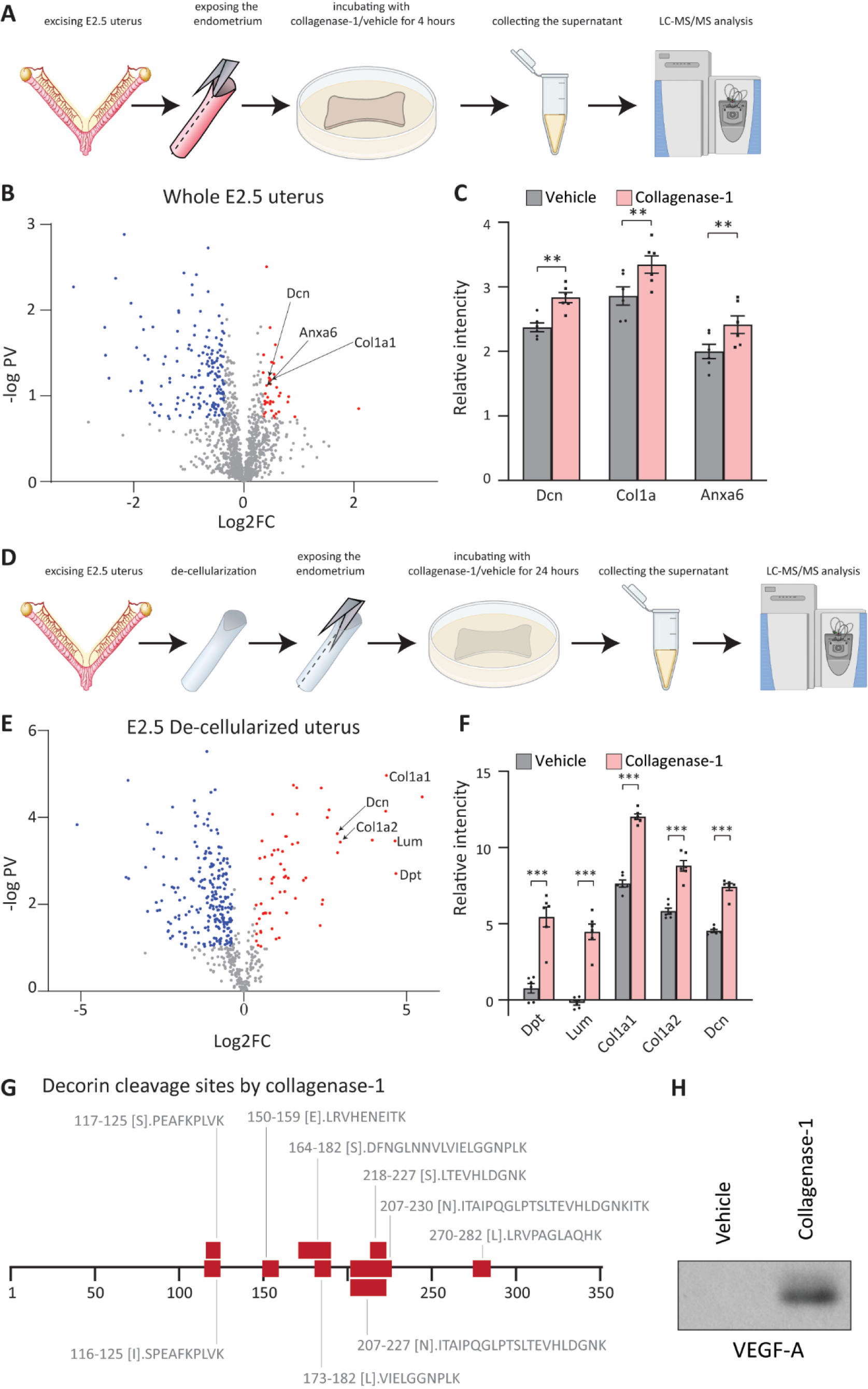
DCN and COL1A1 are collagenase-1 substrates in a pregnant uterus. **(A)** Schematic representation of the experimental set-up: E2.5 uteri were incubated *ex-vivo* with collagenase-1 for 4 hours. The supernatant was subjected to mass spectrometry to reveal potential substrates of collagenase-1 within a pregnant uterus. **(B)** Volcano plot of proteins identified in the supernatant of E2.5 uteri incubated *ex-vivo* with collagenase-1. Red and blue dots are proteins that were upregulated or down regulated (respectively) in the supernatant following incubation with collagenase-1 and pass the threshold of: log2(collagenases-1/vehicle)≥0.4 or ≤-0.4; q≤0.1. **(C)** Bar graph of the matrisome proteins among the upregulated substrates that pass the threshold: collagen alpha 1 (Col1a1, q= 0.057, p=0.0041), decorin (Dcn, q= 0.055, p=0.0038) and annexin A6 (Anxa6 q= 0.062, p=0.0057). ** p<0.01 **(D)** Schematic representation of the experimental set-up: E2.5 uteri were de-cellularized and incubated *ex-vivo* with collagenase-1 for 24 hours. The supernatant was subjected to mass spectrometry to reveal potential substrates of collagenase-1 within the matrix of a pregnant uterus. **(E)** Volcano plot of proteins identified in the supernatant of de-cellularized E2.5 uteri incubated *ex-vivo* with collagenase-1. Red and blue dots are proteins that were upregulated or down regulated (respectively) in the supernatant following incubation with collagenase-1and pass the threshold of: log2(collagenases-1/vehicle)≥0.4 or ≤-0.4; q≤0.1. **(F)** Bar chart showing the most abundant ECM structural proteins and proteoglycan substrates (log2(collagenases-1/vehicle)>2.5) found in E. *** p<0.001 **(G)** Schematic summarizing of semi tryptic decorin peptides detected in the supernatant of E2.5 de-cellularized uteri following incubation with collagenase-1, at respective amino acid positions. **(H)** Representative western blot analysis shows protein expression of VEGF-A in supernatants from decellularized uterus samples that were incubated with collagenase-1 or vehicle for 24 h.

### Vascular permeability and angiogenesis-related processes

Embryo implantation is closely associated with angiogenesis and enhanced uterine vascular permeability, resulting from vessel leakage and the development of new blood vessels^21,22^. Building upon our discovery that VEGF-A is released from a pregnant uterus-ECM following *ex-vivo* exposure to collagenase-1, we delved further to investigate the *in vivo* state of angiogenesis in the tissue after collagenase-1 treatment. We examined various angiogenic makers at two distinct time points after the treatment, E4.5 and E6.5. Remarkably, we observed an increase in the expression of VEGFR-2 at E6.5 and VEGF-A at E4.5 (Fig. 5A, S3). Angiogenesis is endogenously regulated, among other mechanisms, by the balance between the pro-angiogenic VEGF and the anti-angiogenic PEDF factors. A higher VEGF/PEDF ratio promotes angiogenesis, while increased PEDF levels, in particular, impairs embryo implantation^23-25^. In line, we found that coallagenase-1 treatment reduced PEDF levels, concurrent with an increase in VEGF-A (Fig. 5A). Furthermore, to investigate the direct effect of collagenase-1 on PEDF, we conducted an *in vitro* incubation of PEDF protein with collagenase-1 for 24 h, which resulted in a decrease in PEDF levels, indicating its degradation by the enzyme (Fig 5B). Functional assessment to validate the outcome of elevated angiogenic factors was conducted, by staining with CD34, the marker indicating neovascularization^26^. Imaging of the implantation site endometrium revealed higher number of newly formed blood vessels at E6.5 after the WOI in collagenase-1 treated mice (Fig. 5C).

**Fig. 5.**
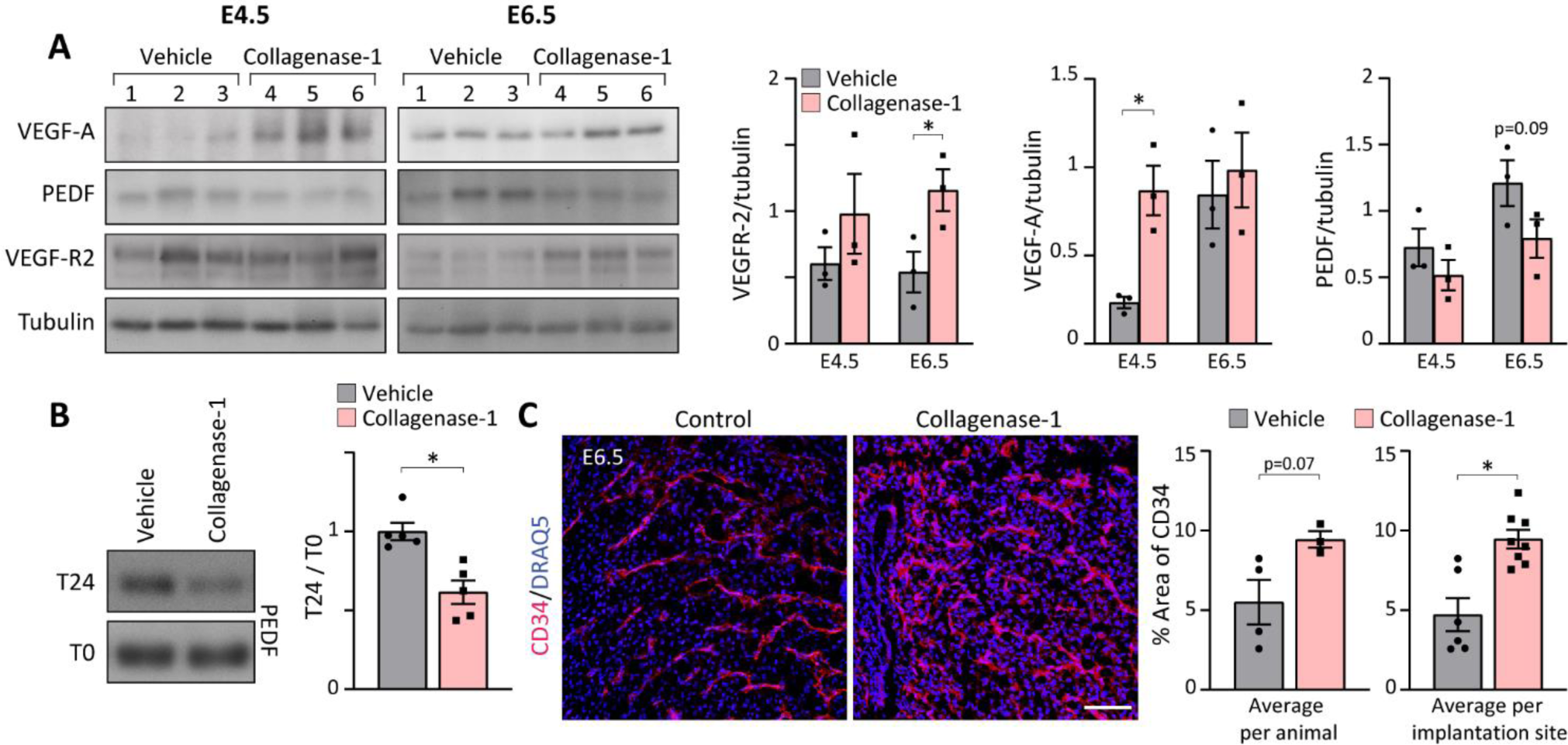
Collagenase-1 promotes vascular permeability at implantation sites. **(A)** Western blot analysis shows the increase of VEGF-R2 and VEGF-A levels in E4.5-E6.5 samples treated with collagenase-1 as well as decrease in PEDF levels (left). Quantification of WB using ImageJ analysis tool (right) (n = 3). **(B)** Representative WB images of incubation of PEDF protein with collagenase-1 *in vitro* for 24 h. Initial PEDF levels at T0 are similar. However, after a 24-hour incubation with collagenase-1, there is a noticeable reduction in PEDF, pointing to its degradation. **(C)** Representative images of immunofluorescent staining against CD34 of E6.5 samples. Covered area quantification was performed using ImageJ analysis tool. Data were analyzed by an unpaired, two-tailed t-test. Results are presented as mean ± SEM, with significance: *p < 0.05.

Further analysis of angiogenesis at the implantation site following collagenase-1 treatment involved the use of magnetic resonance imaging (MRI) at E4.5 to document the dispersion of i.v. injected contrast agents (Fig. S4A). We calculated two key parameters that characterize vascular remodeling and function: permeability surface area (PS) and fractional blood volume (fBV)^22^. This analyses showed that implantation sites in uteri treated with collagenase-1 exhibited slightly elevated values of PS and fBV, although non-significantly, when compared to vehicle-treated uteri (Fig. S4B). These results indicate that collagenase-1 induces a modest increase in uterine vascular permeability without evidence of pathological hypervascularity. Remarkably, immunofluorescent imaging demonstrated that collagenase-1 treatment enhanced penetration and accumulation of the injected contrast agents within the implantation site (Fig. S4C).

Taken together, these results indicate that mild proteolysis of the endometrium promotes local vascular remodeling associated with permeability and angiogenesis, without inducing early pathological hyper-vascularization that could negatively impact pregnancy outcomes.

### NK cell infiltration and upregulation of the cytokine LIF

Intensive collagen degradation has been associated with immune cell migration and infiltration^27,28^. Additionally, embryo implantation involves unique immune activities and pro-inflammatory modulation^29^. Therefore, we examined whether collagen de-tensioning in the uterus affects infiltration of immune cells into the treated endometrium. Flow cytometry analysis of implantation sites at E4.5 showed a specific increase of ∼ 40% in NK cells with no changes in T cells, dendritic cells, or macrophages (Fig. 6A,B, S5A-B). Furthermore, western blot analysis of the NK cell marker (NKp46) at E4.5 confirmed the increase in NK cell infiltration to the implantation site following treatment (Fig. 6C, S5C). These changes induced by collagenase-1 are crucial for improved implantation rates since uterine NK cells are pivotal for successful early pregnancy^30^, and their absence characterizes recurrent implantation failure^31^.

**Fig. 6.**
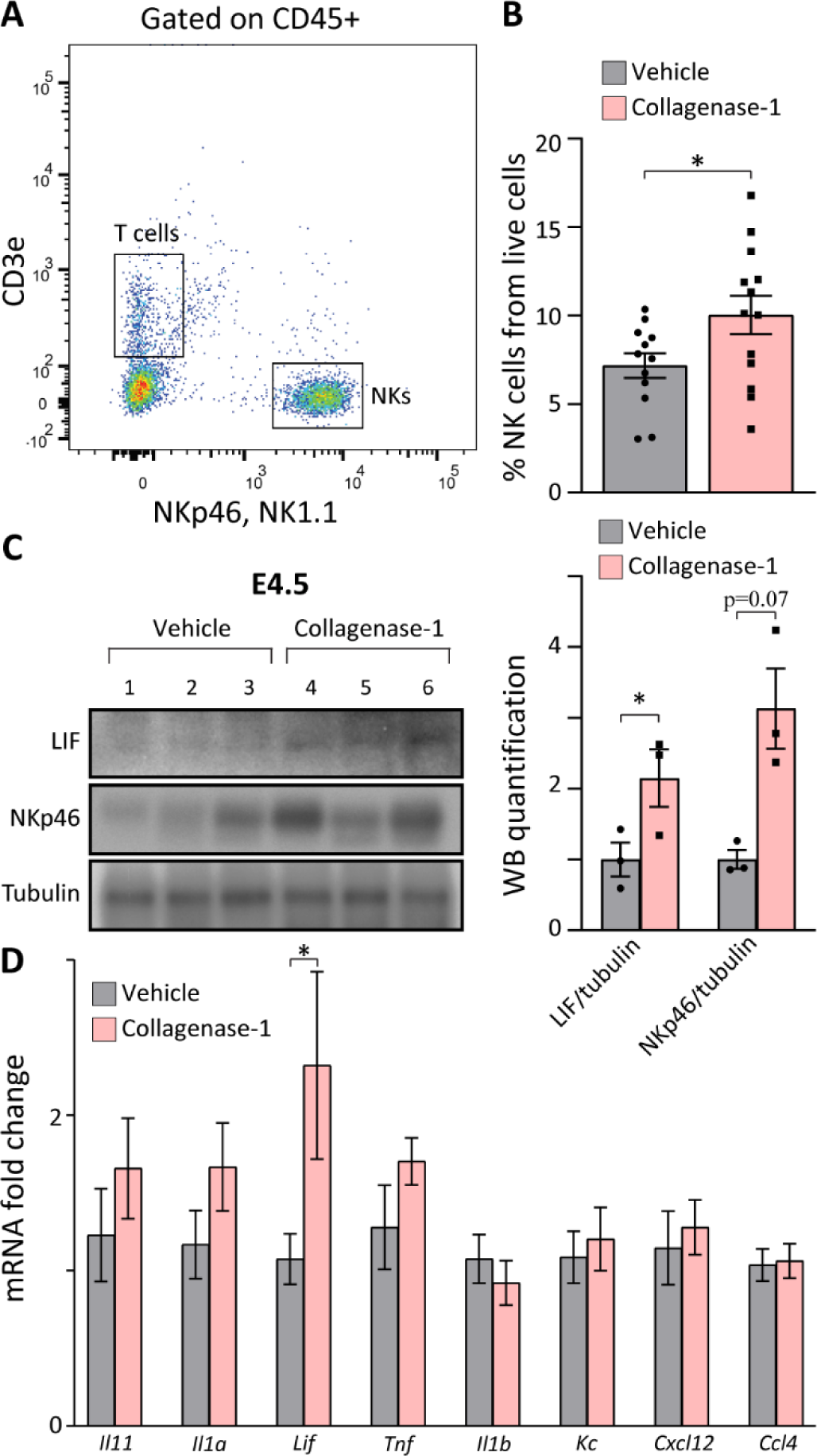
Collagenase-1 treatment increases NK cell infiltration and expression of the cytokine LIF in spontaneous pregnancy model implantation sites. **(A)** A representative flow cytometry image showing the gating strategy used to identify NK cells (CD45+CD3ε−NK1.1+NKp46+). **(B)** Quantification of flow cytometry results showing the percent of NK cells out of live cells at E4.5 (n ≥ 12). **(C)** Representative Western blot (left) and quantification analysis (right) showing expression of LIF and NKp46 in E4.5 implantation sites. Western blot quantification performed using ImageJ analysis tool (n = 3). NKs, Natural killer cells. (**D)** Real-time PCR analysis of cytokines’ transcripts at E4.5 (n = 8). Data was analyzed by an unpaired, two-tailed t-test. Results are presented as mean ± SEM with significance: *p < 0.05.

Since numerous cytokines and chemokines are involved in the embryo-endometrium crosstalk^32,33^, we compared the expression levels of several such molecules at E4.5. We found that collagenase-1 treatment significantly increased the transcript and protein levels of leukemia inhibitor factor (*Lif*) (Fig. 6C,D, S5C), one of the most important pro-inflammatory cytokines involved in successful implantation^34^. Interestingly, there was no significant change in the expression of other tested factors (Fig. 6D), indicating that collagenase-1 treatment did not induce a robust and unbalanced immune response. Of note, establishing a pseudopregnant model (Fig. S6A), we documented the same tissue response to collagenase-1 remodeling, indicated by increased NK cell, LIF and VEGF-A proteins (Fig. S6B-D), demonstrating an embryo-independent process.

Taken together, our results indicate that collagenase-1 treatment promotes a mild yet specific pro-inflammatory signaling in the endometrium, which is recognized as a prerequisite for successful embryo implantation.

### Improved implantation in heat stress and embryo transfer

Next, we investigated whether collagenase-1 treatment could also be effective in models known for exhibiting low embryo implantation rates, i.e., heat stress and embryo transfer (ET). Prolonged exposure to elevated environmental temperature reduces successful pregnancy rates^35-37^. In line, in our experimental model, keeping mice under heat stress conditions of 38°C, we observed reduced embryo implantation rates with an average of only 5 implantation sites per female (Fig. 7A,B). Notably, these females were in a normal and healthy physiological state (Fig. S7A). We hypothesized that the in-uteri structural and signaling changes mediated by collagenase-1 treatment could compensate for the harmful effect of the elevated temperature. Indeed, treatment with collagenase-1 rescued the heat-stress effect as we observed an average of 10.6 implantation sites per female (Fig. 7B).

**Fig. 7.**
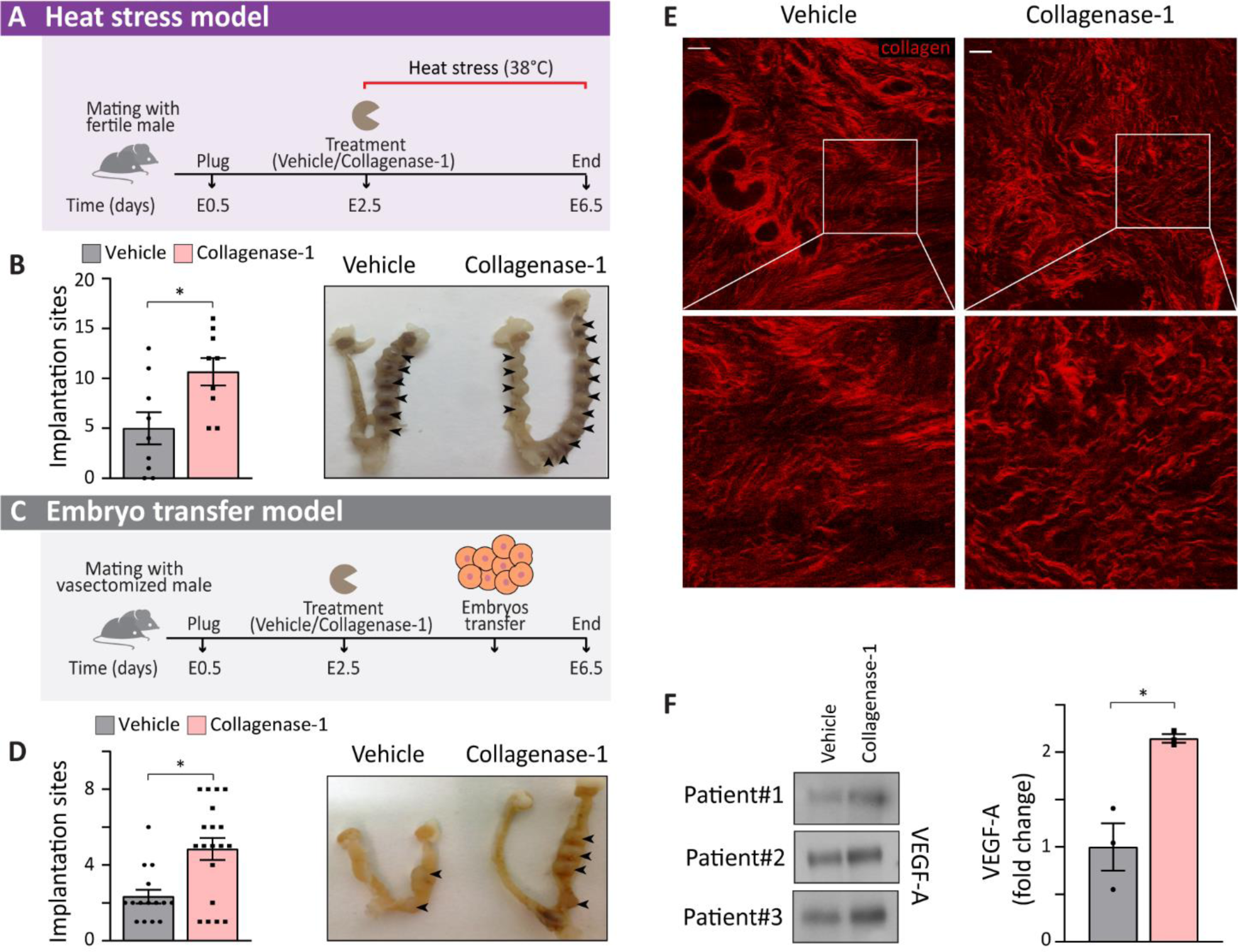
Collagenase-1 improves implantation rate in embryo-transfer and heat-stress models and mechanistically affects human uteri-samples similarly to the mouse tissue. **(A)** A schematic representation of the heat-stress model. **(B)** Quantification of implantation sites and representative images of the uteri at E6.5 in the heat stress model. Arrowheads mark implantation sites (n = 9). **(C)** A schematic representation of the embryo transfer model. **(D)** Quantification of implantation sites and representative images of the uteri at E6.5 in the embryo transfer model. Implantation sites marked by arrowheads (n ≥ 15). **(E)** Representative SHG images of human uterine tissue sections incubated with either 50 nM of collagenase-1 or vehicle for 24 h (scale bar = 50 µm). Enlarged insets presented at the lower panel show misaligned and loosely packed collagen fibers, in the collagenase-1-treated sample. **(F)** Representative Western blot of supernatants from decellularized human uterus samples that were incubated with collagenase-1 or vehicle for 24 h (n=3). Three samples from 3 different women were compared, and for each sample the fold change between collagenase-1 and vehicle was measured. The graph shows the three ratios (p-value=0.09). Quantification of western blot analysis using ImageJ analysis tool. Data was analyzed by an unpaired, two-tailed t-test. Results are presented as mean ± SEM with significance: *p < 0.05.

Another well-known scenario with typical low implantation rates is the embryo transfer model. In humans, *in vitro* fertilization (IVF) and ET show relatively low pregnancy success mainly due to impaired implantation^38,39^. Thus, we investigated whether collagenase-1 treatment can improve implantation rates in a mouse embryo transfer model, in which blastocysts are transferred to a pseudo-pregnant female, which is mated with a vasectomized male (Fig. 7C). A significant two-fold increase in the number of embryo implantations was achieved upon topical administration of recombinant collagenase-1 (Fig. 7D). Furthermore, a similar increase in implantation rates induced by the treatment was observed in a cross-strain embryo transfer protocol, in which cbcF1 embryos were transferred into ICR pseudo-pregnant females (Fig. S7B).

Altogether, these results show a general phenomenon in which collagenase-1-mediated mild remodeling of the endometrium increases the number of implanted embryos regardless of their genetic background. Moreover, this mild and controlled intervention overrides environmental stressors that impair the tightly coordinated embryo implantation process.

### Collagenase-1 similarly affects human endometrial tissue

Finally, a major intriguing question was the feasibility of our approach to create similar effects in assisted reproduction treatment protocols used in human females. To obtain molecular mechanistic indications for the response of the human endometrium to collagenase-1 treatment, we decellularized fresh endometrial biopsies from healthy women. Tissue samples were incubated *ex vivo* with collagenase-1 or vehicle and imaged under two-photon microscopy. Utilizing the SHG imaging modality, we observed collagen fiber de-tensioning in collagenase-1-treated endometrium layers, similar to what was observed in mice (Fig. 7E). Furthermore, we examined the effect of collagenase-1 on the release of matrix-bound VEGF-A. Consistent with the results observed in murine samples, we observed an increased release of 42 kDa dimers of VEGF-A, specifically from the collagenase-1 treated matrices (Fig. 7F).

Figure 8 provides a schematic illustration that summarizes the key factors and molecular mechanisms involved in the processes by which collagenase-1 treatment improves embryo implantation. Overall, our results demonstrate that the mild proteolysis by collagenase-1 generates similar changes in human and mouse tissues in terms of altering the collagen morphology and the release of pro-angiogenic factors. These data indicate the potential use of a specific remodeling enzyme in human reproductive medicine.

**Fig. 8.**
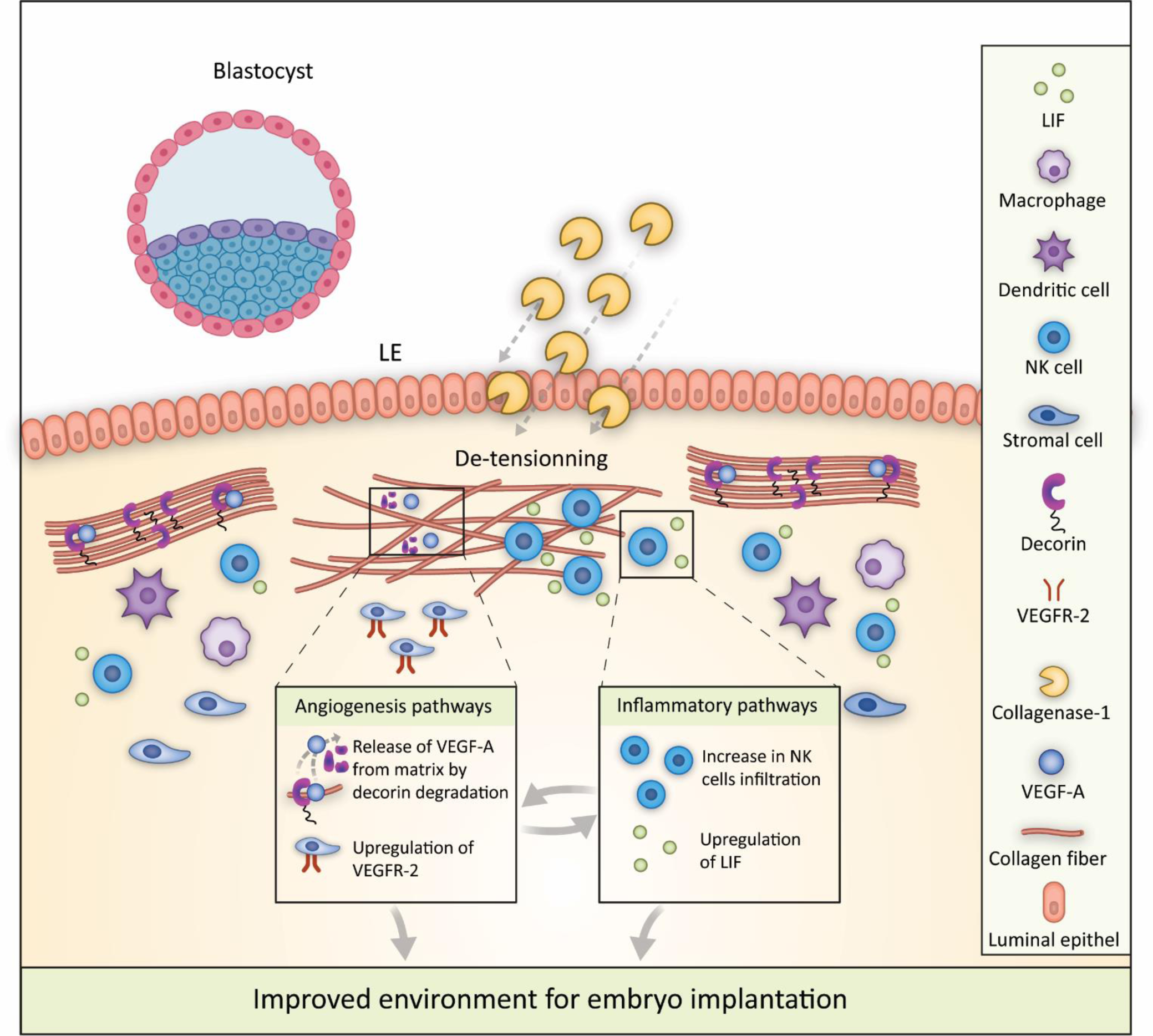
Enhancing embryo implantation: mechanisms of in uteri collagenase-1 treatment. Collagenase-1 administration leads to the de-tensioning of the dense collagen network through mild degradation of collagen and decorin. This process leads a greater accumulation of uterine NK cells secreting LIF. Additionally, collagenase-1 releases VEGF-A from the uterine matrix via decorin proteolysis and upregulates VEGFR-2 expression. These enhancements in inflammatory and angiogenic pathways optimize the receptive environment for embryo implantation.

## Discussion

Embryo implantation is a pivotal event in mammalian reproduction, representing a critical determinant for the establishment of natural early pregnancies and the success of assisted reproductive techniques such as in vitro fertilization–embryo transfer. Given the challenges and limited success rates associated with assisted reproductive technology, extensive research efforts have been dedicated to exploring strategies involving extracellular matrix (ECM) remodeling to improve embryo implantation. Among these strategies, endometrial scratching has emerged as one of the most widely recognized approaches ^40^. Application of this method prior to IVF procedure dramatically increases the chances of implantation by provoking inflammation and endometrial angiogenesis, processes recommended by many clinicians prior to IVF^41-43^. However, the efficiency of this procedure has been questioned by several studies, probably due to reproducibility problems^44-46^. LIF administration has emerged as alternative approach to enhance embryo implantation, since this factor is dysregulated in infertile women experiencing recurrent implantation failure^47^. Unfortunately, LIF treatment did not improve implantation rates and pregnancy outcomes for women with recurrent unexplained implantation failure^48^. Another approach focused on enhancing vascularization. VEGF levels are significantly reduced in uterine fluid during the receptive phase in women with unexplained infertility compared with fertile women^49^. Indeed, mouse embryos cultured with recombinant human VEGF (rhVEGF) had significantly higher implantation rates following blastocyst transfer^50^. However, the net effect of rhVEGF on the uterus environment was not described. Taken together, these studies strongly indicate that factors associated with ECM remodeling have the potential to improve embryo implantation. However, a deeper understanding of ECM remodeling mechanisms and their role in embryo implantation is required to better harness this strategy for medical reproductive interventions.

Our findings unequivocally demonstrate that the judicious application of collagenase-1 induces a mild, yet highly targeted extracellular matrix (ECM) remodeling process, which orchestrates crucial physiological changes within the uterine microenvironment to foster optimal conditions for successful embryo implantation. In particular, collagenase-1 actively participates in the intrinsic and dynamic remodeling of vital ECM components essential for successful implantation, namely fibrillar collagens and decorin. Notably, the degradation of these pivotal proteins mirrors the natural course of events during peri-implantation and the early stages of implantation^51-53^, indicating that collagenase-1 administration must be tightly synchronized before embryo attachment to the uterus wall. Physiological de-tensioning of endometrial collagen fibers associates with trophoblast proliferation and invasion^8^ as well as angiogenesis^19^. We found that facilitating this process exerts a slight increase in permeability surface area of blood vessels and fractional blood volume. Enhanced angiogenesis is evident through the upregulation of VEGF-A and VEGFR2 expression, the release of matrix-bound VEGF-A, and reduced levels of the angiogenic suppressor PEDF. Collagenase-1 has been previously shown to promote VEGFR-2 expression through the stimulation of protease activated receptor-1 (PAR-1)^54^. The release of matrix-bound VEGF-A primarily occurs through the degradation of proteoglycans, with our proteomics results pointing to decorin degradation as its main source. Remarkably, collagen de-tensioning creates space for increased infiltration of NK cells. NK cells play a significant role in trophoblast invasion, vascularization, and the initiation of decidualization during embryo implantation^55^. Interestingly, among tested embryo implantation-related cytokines, only LIF showed elevated gene expression and protein levels. The increase of both, LIF and NK cells infiltration at E4.5 indicates that these cells are the source of LIF. LIF affects trophoblast invasion, adhesion, trophoblast cell proliferation and may stimulate their adhesion to the ECM via different signaling pathways^56-58^.

Using a pseudo-pregnancy model at E4.5, we observed that collagenase-1 induces pro-inflammatory and angiogenesis pathways independently of the presence of embryos (Fig. S7). These data further confirm that collagenase-1 primes the uterine environment preparing the embryo-endometrium interface for successful implantation. Clinically significant, collagenase-1 administration was able to rescue low implantation-rates in the heat-stress and embryo transfer models, without compromising endometrial integrity or normal offsprings development. Crucially, a feasibility test of collagenase-1 application on human endometrial samples ex-vivo showed induction of spatial re-organization of collagen fibers and the release of VEGF-A, mechanisms similar to those characterized in mice. A schematic model illustrating the mechanisms by which enforced collagenase-1 remodeling facilitates embryo implantation is presented in Fig. 8. In summary, our study reveals that the administration of collagenase-1 directly into the uterine environment represents a powerful strategy to augment endometrial receptivity, consequently enhancing the chances of successful embryo implantation. The key mechanisms underlying this enhancement lie in collagen de-tensioning, a process triggered by collagenase-1, and the concurrent induction of angiogenesis within the endometrial tissue. Accordingly, our study not only sheds light on the fundamental biology of embryo implantation but also offers compelling implications for the field of biomedical engineering.

The application of collagenase-1 as a localized treatment holds immense promise across diverse domains. In the realm of livestock breeding, where efficient reproductive outcomes are crucial for agricultural productivity and environmental sustainability. Moreover, the implications of our study extend to the realm of assisted human reproductive protocols, encompassing both natural conception and in vitro fertilization (IVF)-based approaches. Collagenase-1 emerges as a biomedical tool, capable of fine-tuning the uterine microenvironment to create a receptive milieu for embryonic attachment. This innovation has the potential to revolutionize the success rates of IVF procedures. This convergence of biological insights and biomedical engineering applications represents a significant step towards addressing pressing challenges in reproduction, with far-reaching implications for both agriculture and healthcare.

## Materials and Methods

### Collagenase-1 preparation, activation and enzymatic assays

Collagenase-1 was overexpressed and purified as described previously^16^. Before each experiment, collagenase-1 was activated with 1 mM APMA (4-aminophenylmercuric acetate) in TNC buffer (50 mM Tris (pH 7.5), 150 mM NaCl, 10 mM CaCl2) at 37 °C for 60 min. The solution with activated collagenase-1 was then fractionized by centrifugation (5 min, 400*g*). Only the upper fraction of the enzyme was used in all others experiments. The enzymatic activity of collagenase-1 was measured at 37 °C by monitoring the hydrolysis of the fluorogenic peptide Mca-Pro-Leu-Gly-LeuDpa-Ala-Arg-NH2 at λex = 340 nm and λem = 390 nm as previously described^59^.

### Seeding blastocysts on fascicle-derived ECM

Fascicle-derived ECM was prepared from the tails of adult Norwegian rats (6 month). Rat tails were dissected, and tendon fascicles (diameter ∼0.6 mm, length ∼7 mm) were gently extracted and were washed extensively in TNC buffer (50 mM Tris (pH 7.5), 150 mM NaCl, 10 mM CaCl_2_) to remove the macroscopic tissue debris and proteases remains. ECM samples were incubated with 500 nM collagenase-1 or vehicle (TNC) at 30 °C for 24 h^16^. After incubation, tails were extensively washed with PBS. Five mouse blastocysts were seeded on top of the ECMs placed in 24-well cell culture plate for 4h in 37°C and then imaged by EVOS M5000.

### Scanning electron microscopy (SEM)

Fascicle-derived native collagen-rich ECM or E2.5 decellularized mouse uterus tissues (longitudinal cut) were fixed in a 0.1-M cacodylate buffer solution (pH 7.4) containing 2.5% paraformaldehyde and 2.5% glutaraldehyde (pH 7.2) for 60 min at room temperature and then were washed three times in the same buffer. The samples then were stained with 4% (wt/vol) sodium silicotungstate (Agar Scientific, UK) (pH 7.0) for 45 min with following dehydration through an ascending series of ethanol concentrations up to 100% ethanol. Next, the samples were dried in a critical point dryer (CPD) and attached to a carbon sticker. Finally, the samples were coated with thin gold/ palladium (Au/Pd) layer. The samples were observed under a Zeiss FEG Ultra55 SEM operating at 2 kV.

### Mice

All animals were obtained from Envigo Laboratories (Jerusalem, IL). On arrival, male mice were housed individually, and female mice were housed 3 to 5 per cage in animal rooms maintained at 20 to 22 °C with an average relative humidity of 35% under a 12:12 h light: dark cycle and were housed in standardized ventilated microisolation caging. All experiments and procedures were approved by the Weizmann Institute of Science Animal Care and Use Committee (IACUC approval no. 27170516-2).

### Topical administration of collagenase-1 or vehicle in a spontaneous pregnancy protocol

Female ICR or C57BL/6J mice (8-10 weeks) were copulated with fertile ICR or C57B/6J male mice (8-12 weeks), respectively. Females which presented a vaginal plug on E0.5, were administered with l.5 µL of 15.5 µM activated recombinant collagenase-1 (total amount 1.25 µg) or 1.5 µL of vehicle as control (TNC buffer) at E2.5 using the NSET™ Device of ParaTechs (Kentucky, US): mice were placed on a wire-top cage and allowed to grip the bars. The small and large specula were placed sequentially into the vagina to open and expose the cervix. Then, the NSET catheter was inserted through the large speculum, past the cervical opening, and into the uterine horn allowing the topical administration of recombinant collagenase-1 or vehicle. Uteri were excised 1 h, 2 days or 4 days after treatment (E2.5, E4.5 and E6.5 respectively). On E4.5 and E6.5 the number of implantation sites was counted. On E6.5 implantation sites were visible and on E4.5 Evens-blue dye was injected intravenously 10 min before euthanizing which allows the visualization of the implantation sites. For generating fluorescent venus embryos, C57BL/6J female mice were mated with Myr-Venus homozygote males ^60^. <u>In the heat-stress protocol</u>: After treatment at E2.5, mice were transferred to preheated housing cages at 38 °C for 4 days and were sacrificed at E6.5. Selected females were injected with a thermal microchip in the back of the mice a week before the experiment. The microchip injection was done under an isoflurane sedation coupled with Carprofen (5 mg/kg) as an analgesic.

### Administration of collagenase-1 or vehicle in a non-surgical embryo transfer protocol

ICR mice (8-10 weeks) were mated with vasectomized male to achieve a pseudo-pregnant state. Females which presented a vaginal plug on E0.5, were administered at E2.5 with 1.25 µg activated recombinant collagenase-1 or vehicle as control using the NSET™ Device as describe above. After 15 min, embryos at the blastocyst stage were transferred also through the NSET catheter into the uterus (10 embryos per mice). Then, the device and specula were removed, and the mouse was returned to its home cage. The number of implanted embryos were counted and recorded on E6.5.

### H&E staining of frozen sections

E4.5 uteri samples treated with collagenase-1 or vehicle embedded in OCT were cross-sectioned (12μm) on glass microscope slides. Slides were washed with phosphate buffered saline (PBS) before staining. Then, slides were incubated in Hematoxylin for 3 min and washed with tap water. After, slides were dipped in 95% ethanol and incubated in Eosin for 45s. Next, samples were dipped in 95% ethanol and incubated 2 min in 100% ethanol. Finally, samples were incubated in xylene for 2 min and were mounted in mounting medium.

### Two-photon microscopy and second-harmonic generation (SHG)

Snap-frozen murine uterine samples (excised at E2.5 and E4.5) were thawed in PBS, cut longitudinally, place on a slide as the endometrium facing up and covered with cover slip. Then samples were imaged using a two-photon microscope in a SHG mode (2PM: Zeiss LSM 510 META NLO; equipped with a broadband Mai Tai-HP-femtosecond single box tunable Ti-sapphire oscillator, with automated broadband wavelength tuning 700–1020 nm from Spectraphysics, for two-photon excitation). For second-harmonics imaging of collagen, a wavelength of 800-820 nm was used (detection at 390-450 nm). For imaging the myometrium layer and venus-embryos uteri samples embedded in OCT were cross-sectioned (50μm) on glass microscope slides.

### Entropy analysis of the distribution of fiber orientations

Detection of collagen fibrils of vehicle and collagenase-1 treated endometrium samples was performed using ImageJ^61^. A mask highlighting collagen fibers was generated using the Tubeness plugin, and was followed by orientation analysis using the Directionality plugin^62^. This gave rise to plots and spreadsheets of distribution of fiber orientations throughout the stacked images (supplementary macrocode file named “Orientation”). Next, the orientation data was analyzed using Matlab. The degree of orientation diversity, i.e., distribution dispersion was measured by calculating the edge-orientation entropy that refers to the probability of encountering particular orientations in an image^63^, and according to the Shannon index^64^. For every orientation i, let *p*_*i*_ be the probability of the occurrence of orientation *i* in the image. Entropy or Shannon index (H’) of the probability distribution is defined as *H*^′^ = 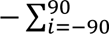 *p*_*i*_ ln *p*_*i*_ (supplementary Matlab code file named “Entropy”). On E2.5 we analyzed 3 vehicle-treated females (4-12 images per each) and 5 collageanse-1-treated females (3-10 images per each). On E4.5 we analyzed 3 vehicle-treated females (4-16 images per each) and 3 collageanse-1-treated females (2-15 images per each).

### Fibers density analysis

Fiber’s density analysis was done by ImageJ software: For endometrium analysis a z-projection of max intensity was applied for 13 µm of the endometrium layer and only fibrillar areas were chosen. Next, auto threshold on the default setting was applied, and Analyze Particles plugin was used to detect the collagen-covered area. Size of particles was chosen as 0.172-infinity. On E2.5, we analyzed 3 vehicle-treated females (4-12 images per each) and 5 collageanse-1-treated females (3-10 images per each). On E4.5, we analyzed 3 vehicle-treated females (4-16 images per each) and 3 collageanse-1-treated females (2-15 images per each). For myometrium analysis, a single section was taken and the myometrium area was chosen for auto threshold and analyze particles.

### De-cellularization of uterine tissue

Mice (E2.5) or woman biopsy uteri samples treated with collagenase-1 or vehicle were incubated in a de-cell solution containing 3% Triton-100 (6 h, 4 °C) and then in a de-cell solution with 0.4% Triton-100 overnight at 4°C [de-cell solution: 1.5 M NaCl, 50 mM Tris pH 8, 50 mM EDTA, protease inhibitor cocktail (Roche)]. Samples were washed three times in ddH_2_O and then incubated with 0.5% sodium deoxycholate (60 min, 25 °C) to remove lipid remaining. Samples were washed again three times in ddH_2_O and stored at 4 °C until use.

### SBF-SEM

Samples were prepared based on methods described previously^18^. In brief, de-cellularized E2.5 uteri samples treated with collagenase-1 or vehicle were fixed in situ by using 2% (wt/vol) glutaraldehyde in 0.1 M phosphate buffer (pH 7), en-bloc stained in 2% (wt/vol) osmium tetroxide, 1.5% (wt/vol) potassium ferrocyanide in 0.1 M cacodylate buffer (pH 7.2), for 1h at 25 °C. Specimens were then washed again 5 x 3 min in ddH_2_O and then specimens were placed in 2% osmium tetroxide in ddH_2_0 for 40 min at 25 °C. Samples were then infiltrated and embedded in TAAB 812 HARD following sectioning using a Gatan 3View microtome within an FEI Quanta 250 SEM. For endometrium samples, a 41 μm × 41 μm field of view was chosen and imaged by using a 4096 × 4096 scan, section thickness was set to 100 nm in the Z (cutting) direction. Z volumes datasets comprised 700 images (100 μm z depth). The IMOD suite of image analysis software was used to build image stacks, reduce imaging noise, and generate 3D reconstructions. Fibrils were contoured using functions in the IMOD image analysis suite.

### Proteomic analysis

Mice uteri samples (E2.5) were excised and a vertical incision was made to gain maximum exposure of the endometrium. The samples were incubated in 100 µL of DMEM supplemented with TNC (vehicle) or 100nM collagenase-1 for 4 h at 37°C. For matrisome proteins enrichment, the E2.5 were de-cellularized, then a vertical incision was made and the samples were incubated in 100 µL of TNC buffer supplemented with 100 nM collagenase-1 for 24 h at 37°C (for control no enzyme was added). At the end of incubation, the supernatants were collected. 10µg total protein from each sample was mixed 1:1 with 8M urea, 100mM Tris/HCL (pH8.5) to a final concentration of 4M urea. TCEP was added to a final concentration of 10mM and incubated in shaking at RT for 30 min. Then, CAA was added to a final concentration of 40mM and incubated in shaking at RT for 30 min. The samples was then diluted 1:3 with 50 mM ammonium bicarbonate buffer. In-solution digestion was performed with LysC-Trypsin mix (1:100 enzyme: protein ratio) and trypsin (Promega; 1:50 enzyme: protein ratio). Peptides were desalted on C_18_ stage tips, vacuum dried, and resuspended in 0.1% TFA.

Peptides were introduced into the mass spectrometer by means of Waters nanoAcquity HPLC system connected to a Symmetry trap column [180umX20mm] and Analytical column HSS T3 75umX250mm, both from Waters. Data was acquired on Q exactive HF mass spectrometer (Thermo Scientific) with Top 15 method. Raw files were searched against the Mus musculus [Taxonomy ID: 10090] database compiled from the UniProt reference proteome using Sequest from Proteome Discoverer™ 2.4 software (Thermo). The following parameters were selected for database searches: semi-Trypsin for enzyme specificity with tolerance of one missed cleavage; carbamidomethyl(C) as fixed modifications, and acetyl(N-term), pyroQ (N-term), oxidation(M), deamidation (NQ), as variable modifications. Precursor mass error tolerance of 10 ppm and fragment mass error at 0.02 Da. Percolator was used for decoy control and FDR estimation (0.01 high confidence peptides, 0.05 medium confidence).

### Cell extraction and western blotting

Frozen uterus tissues of 1 cm length were washed in PBS, homogenized in 0.5mL RIPA buffer (EMD Millipore, Burlington, MA, USA) with a protease inhibitor (Roche, Basel, Switzerland) using a hand homogenizer and centrifuged (14000 x*g*, 15 min, 4 °C). Supernatants were resuspended in sample buffer (200 mM Tris pH 6.8, 40% glycerol, 8% sodium dodecyl sulfate (SDS), 100 mM dithiothreitol (DTT), 0.2% bromophenol blue), and boiled for 5 min. Tissue extracts were then subjected to SDS polyacrylamide gel electrophoresis (PAGE) and transferred onto nitrocellulose membranes (Whatman, PA, USA) by electro-blotting. Membranes were blocked in Tris-buffered saline with Tween 20 (TBST) buffer (200 mM Tris pH 7.5, 1.5 M NaCl, 0.5% Tween 20) and 2% bovine serum albumin (BSA, 60 min, 25°C) and then incubated with the corresponding primary Ab (60 min, 25 °C), washed three times with TBST and incubated with horseradish peroxidase (HRP)-conjugated secondary antibody (60 min, 25°C). Quantification of the band intensities was performed using ImageJ analysis tool.

Antibodies used in this study: VEGF (A-20): sc-152 (Santa Cruz Biotechnology), LIF (R&D systems, AF449), VEGF-R2 (cell signaling, 55B11), NKp46 (R&D systems, AF2225), β-tubulin (Santa cruz, sc-9104). Polyclonal anti-PEDF was kindly provided by Dr. Galia Maik-Rachline (Prof. Roni Seger lab, Weizmann institute, Rehovot, Israel). Secondary antibodies (both anti-rabbit and mouse) conjugated to horseradish peroxidase (HRP) were purchased from Jackson ImmunoResearch (cat No.111-001-003 and 115-001-003, respectively). Antibodies were used at the manufacturer’s recommended dilution.

### Immunofluorescence staining

Uterine samples were fixed with PBS 4% PFA, paraffin embedded and sectioned (4 μM). Sections were deparaffinized and epitope retrieval was performed in citric acid buffer (pH 6). Samples were blocked in PBS, 20% normal horse serum, and 0.2% Triton X-100 and then incubated with primary Ab in PBS, containing 2% normal horse serum and 0.2% Triton X-100 (60 min, 25 °C). Next, samples were washed three times in PBS and incubated with a secondary antibody (60 min, 25 °C) and mounted in a mounting medium. Primary antibodies: CD34 (CL8927PE; Cedarlane, Ontario, Canada). For each image, a suitable threshold was applied for CD34 channel, and Analyze Particles plugin was used to detect the covered area.

### MRI imaging

MRI experiments were performed at 9.4 T on a horizontal-bore Biospec spectrometer (Bruker) using a linear coil for excitation and detection (Bruker) as reported previously^22^. The animals were anesthetized with isoflurane (3% for induction, 1%–2% for maintenance; Abbott Laboratories) in 1 L/min oxygen, delivered through a muzzle mask. Respiration was monitored, and body temperature was maintained using a heated bed. The pregnant mice were serially scanned at E4.5. Three-dimensional gradient echo (3D-GE) images of the implantation sites were acquired before, and sequentially, for 30 min after i.v. administration of the contrast agent. A series of variable flip angle, precontrast T1-weighted 3D-GE images were acquired to determine the precontrast R1 (repetition time [TR]: 10 msec; echo time [TE]: 2.8 msec; flip angles 5°, 15°, 30°, 50°, 70°; 2 averages; matrix, 256 **׳** 256 **׳** 64; field of view [FOV], 35 **׳** 35 **׳** 35 mm^3^). Postcontrast images were obtained with a single flip angle (15°). During MRI experiments, the macromolecular contrast agent biotin-BSA-GdDTPA (80 kDa; Symo-Chem), 10 mg/mouse in 0.2 mL of PBS, was injected i.v. through a preplaced silicone catheter inserted into the tail vein. The MRI scans allowed quantification of the fBV and the permeability surface area product (PS) of embryo implantation sites, as previously reported^22^. In brief, the change in the concentration of the administered biotin-BSA-GdDTPA over time (Ct), in the region of interest, was divided by its concentration in the blood (Cblood); calculated in the region of interest depicting the vena cava, also acquired during MRI, and extrapolated to time 0). Linear regression of these temporal changes in Ct/Cblood yielded 2 parameters that characterize vascular development and function: (a) fBV (fBV = C0/Cblood), which describes blood-vessel density and is derived from the extrapolated concentration of the contrast agent in implantation sites, at time zero, divided by the measured concentration in the vena cava, approximately 5 min after i.v administration, and (b) PS = ([Ct – C0]/[Cblood **׳** t]), which represents the rate of contrast agent extravasation from blood vessels and its accumulation in the interstitial space and which is derived from the slope of the linear regression of the first 15 minutes after contrast agent administration (t = 15). Mean fBV and PS were calculated separately for single implantation sites, considering homogeneity of variances between mice. At the end of the MRI session, embryo implantation sites were harvested and immediately placed in 4% PFA after sacrificing the pregnant mice by cervical dislocation. Slides were then washed in PBS and incubated in Cy3 or Cy2-conjugated streptAvidin (Jackson Immunoresearch Laboratories, PA, USA), diluted 1:150 in PBS for 45 min.

### Quantitative Real Time PCR (qRT-PCR)

E4.5 uteri samples treated with collagenase-1 or vehicle were homogenized using a hand homogenizer. Total RNA was isolated using PerfectPure RNA Tissue Kit (5 Prime GmbH, Deutschland). 1 μg of total RNA was reverse transcribed using High Capacity cDNA Kit (Applied Biosystems Inc. MA, USA). qRT-PCR was performed using specific primers with SYBR Green PCR Master Mix (Applied Biosystems Inc.) on ABI 7300 instrument (Applied Biosystems) readouts were normalized to a B_2_M housekeeping. Primer sequences are listed in Table 1 below. Data are presented as mean fold change using the 2^−ΔΔ*CT*^ method^65^. The standard error of the mean (SEM) was calculated on the 2^−ΔΔ*CT*^ data, as was the statistical analysis.

**TABLE 1.**
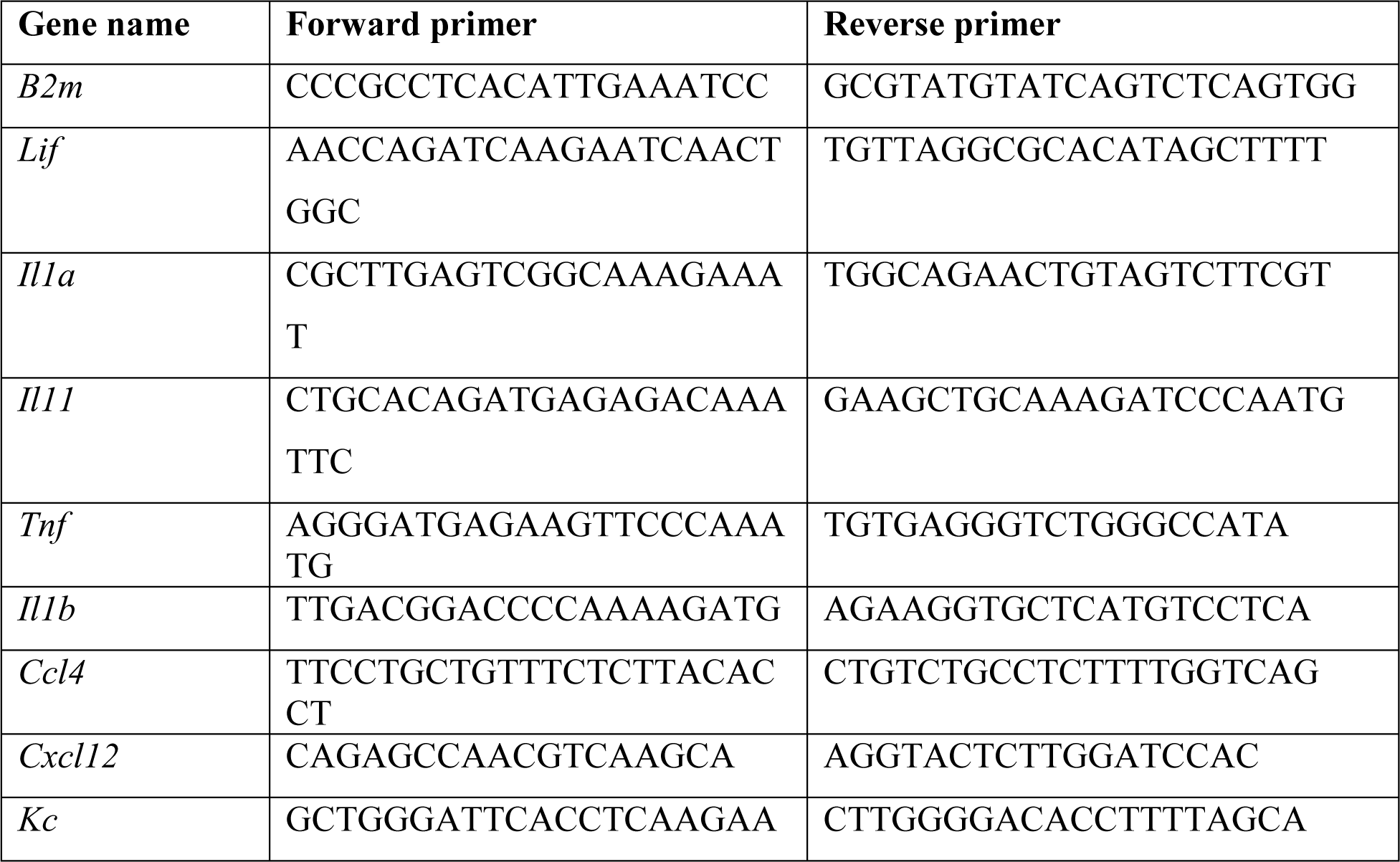
Primer sequences used for qRT-PCR.

### Cell isolation from uterus tissue

Single implantation sites were stained using I.V injection of Evan’s blue dye (Sigma-Aldrich, Rehovot, Israel), excised and isolated on E4.5. Tissues were minced into small fragments and incubated in shaking for 40 min at 37 °C with PBS +/+ containing 0.5 mg/mL collagenase type IV (Sigma-Aldrich, Rehovot, Israel) and 0.1 mg/mL DNase I (Roche). Digested tissue was filtered and smashed with a syringe plunger through a 250 μm nylon sieve in FACS buffer [PBS, 2% fetal calf serum (FCS), 2 mM ethylenediaminetetraacetic acid (EDTA)] to mechanically dissociate the remaining tissue. The supernatant cell pellet was then centrifuged at 390 x*g*, and the pellet cells were lysed for erythrocytes using a red blood cell lysis buffer (Sigma-Aldrich, Rehovot, Israel) (2 min, 25 °C).

### Flow cytometry analysis

The following anti-mouse antibodies were used: CD45 (30-F11), CD3ε (clone 145-2C11), NKp46 (clone 29A1.4), NK1.1 (clone PK136), CD64 (clone 10.1), CD11b (clone M1/70), CD11c (clone N418), IAb (clone AF6-120.1) - all purchased from BioLegend (San Diego, USA). Anti-mouse F4/80 (clone A3-1) was purchased from BIORAD. The cells were incubated with the antibodies for 30 min in FACS buffer (dark, 4 °C) and then washed once with FACS buffer. Cells were analyzed with BD LSR II, special order system (BD Biosciences). Flow cytometry analysis was performed using FlowJo software (TreeStar, Ashland, OR, USA).

### Human uterine samples

Fresh uterine samples from healthy women biopsies were obtained by Dr. Eitan Ram, Gynecologic Oncology Division, Helen Schneider Hospital for Women, Rabin Medical Center; Petah-Tikva. Helsinki approval no. 0450-16-RMC. The tissues were decellularized using the same procedure already described. The samples were treated by vehicle or 50 nM collagenase-1 in a volume of 50 uL at 37 °C for 8 h.

### Statistical Analysis

Statistical analyses were carried out using GraphPad Prism software (VIII, GraphPad Software Inc., La Jolla, CA). Data were analyzed by unpaired, two-tailed t-test to compare between two groups. Multiple comparisons were analyzed by one-way analysis of variance (ANOVA). After the null hypothesis was rejected (p < 0.05), Tukey’s Honestly Significant Difference or Dunnett tests were used for follow-up pairwise comparison of groups in the one-way ANOVA. Data are presented as mean ± SEM in the figures; values of p < 0.05 were considered statistically significant (∗P < 0.05, ∗∗P < 0.01, ∗∗∗P < 0.001).

## Data and materials availability

All data needed to evaluate the conclusions in the paper are available in the main text or the supplementary materials.

## Author contributions

Conceptualization: EZ, ISo, ISa. Methodology: EZ, TG, ES, RH, OG, GM, KEK, MN, ND. Software: GM, ES, OG, EZ, TG. Validation: EZ, TG, RH, IA, ISo, OK, ISa. Formal analysis: EZ, TG, ISo, RH, IA, OG, GM. Investigation: EZ, TG, IA, KEK, ISo, Isa. Resources: RE, KEK, MN. Writing - original draft: ISo, EZ, TG, ISa. Writing—review & editing: ISo, TG, EZ, IA, KEK, ES; RH, GM, OG, OK, MN, ND, ISa. Visualization: EZ, TG, ISo, ISa. Supervision: ISo, ISa. Funding acquisition: ISa.

## Competing interests

“Compositions for remodeling extracellular matrix and methods of use thereof” (U.S. patent no. 10722560). Assignee: NanoCell Ltd, Investors: ISa, ISo, EZ. All other authors declare that they have no competing interests.

## Supporting information

Macrocode file, Orientation.

Macrocode file, Entropy.

## List of Abbreviations

BM: basement membrane
ECM: extracellular matrix
ET: embryo transfer
fBV: fractional blood volume
GFP: green fluorescent protein
IVF: *in vitro* fertilization
LIF: leukemia inhibitor factor
MMP: matrix metalloproteinase
MRI: magnetic resonance imaging
PS: permeability surface area
SBF-SEM: serial block-face scanning electron microscope
SEM: scanning electron microscopy
SHG: second-harmonic generation
VEGFA: vascular endothelial growth factor A
VEGFR2: vascular endothelial growth factor receptor 2
WOI: window of implantation

## Acknowledgments

We thank Dr. Alina Berkovitz for embryo preparations and fruitful discussions, Ms. Anna Aloshin for collagenase-1 purification, and Ms. Yinhui Lu at the University of Manchester for collecting SBF-SEM images. We thank Dr. Ori Brenner for the assistance in histopathological characterization of mice uteri. The images in this paper were acquired at the Optical Imaging & Translational Bioengineering Unit, Department of Veterinary Resources, and at the Advanced Optic Imaging Unit, de Picciotto-Lesser Cell Observatory In memory of Wolfgang and Ruth Lesser at the Moross Integrated Cancer Center Life Science Core Facilities, Weizmann Institute of Science. We thank Tevie Mehlman for the MS data acquisition. This research was supported by the Israeli Science Foundation (1226/13. I.Sa.).

## Notes

### Summary of Updates

New Figure 4 (old Fig. 4 moved to supp materials). New data in Figure 5. Statistics analyses.

