## Supplementary material for "Enhancing Uterine Receptivity for Embryo Implantation through Controlled Collagenase Intervention": Macrocode file, Orientation.: Orientation.ijm

```
// Create Directionality Stack
// Ofra Golani, Guillaume Molodij 16/10/2017
var CollageCh = 4;		// channel of interest 
var tubenessSigma = 1;	// sigma is used both for tubeness and OrientationJ, it is based on the evaluation of the maximum of coherence determined by
						// Orientation J Dominant Direction 
var ResultsSubFolder = "Results";
var fileExtention = ".tif";

directory = getDirectory("Open Image folders");
resFolder = directory + ResultsSubFolder + File.separator; 
File.makeDirectory(resFolder);
print("inDir=",directory," outDir=",resFolder);

setBatchMode(true);
ProcessFiles(directory, resFolder);


function ProcessFiles(directory, resFolder) {
	fileListArray = getFileList(directory);
	for (fileIndex = 0; fileIndex < lengthOf(fileListArray); fileIndex++) {
		if (endsWith(fileListArray[fileIndex], fileExtention)) {
			open(directory+fileListArray[fileIndex]);	
			ProcessFile(directory, resFolder);
			run("Close All");
		} 
	} 
} 

function ProcessFile(directory, resFolder) {
	y=getHeight;
	x=getWidth;
	z=nSlices;
	print(x,y,z);
	imName=getTitle();
	// extract only the channel of interest
	selectWindow(imName);
	run("Duplicate...", "title=Collagen duplicate channels="+CollageCh);
	origStack=getTitle();
	for(i=0; i<nSlices; i++){
		num=toString(i+1);
		selectWindow(origStack);
		if (nSlices >2) {
			Stack.setSlice(i);	
		}
		print(i);
		run("Duplicate...", "title=curSlice");
		run("Tubeness", "sigma="+tubenessSigma+" use");
		//waitForUser
		selectWindow("tubeness of curSlice");
		rename("tubenessImg");
		tubenessName=getTitle();
    	run("Directionality", "method=[Local gradient orientation] nbins=90 histogram=-90 display_table");
		selectWindow("Directionality histograms for tubenessImg (using Local gradient orientation)");
    	name = imName+num;
    	name = name + ".csv";
    	print(resFolder+name); 
    	saveAs("Results", resFolder+name);
    	//waitForUser;
  		selectWindow(name);
  		run("Close");
  		close("tubenessImg");
  		close("curSlice");
	}
	run("Close All");
}
```
