## Supplementary material for "Enhancing Uterine Receptivity for Embryo Implantation through Controlled Collagenase Intervention": Macrocode file, Entropy.: Entropy.m

```
% Capturing the disorientation effect of control and treated samples

% Method: evaluate the level of disorientation by measuring the entropy of
% the oritentation distribution for each slice of each sample (in the form
% of a z-stack). The mean entropy is calculated per image, then per mouse, as well as
% the mean of all control or all treated samples. A plot of all per mouse
% entropies(colored points)and total means (black crosses) is created in Fig. 1,
% comparing control vs. treated. The p-value for a two-sample t-test is
% reported. If the removal of outliers is required, Fig. 2 will display the
% plot without outliers and a new p-value (pNO) will be reoprted. Outliers
% are defined as being 2 or 3 (according to the value of the variable
% cutOff) mean absolute deviations (MADs) from the median of their group.
% If cutoff is chosen to be changed, a new Fig. 3 will be created with the
% new cutoff.
% Outliers are removed within each mouse. Each mouse should have all its
% images in one folder.

% Use first Fiji macro Orientation.ijm to create the .csv tables and set the result
% directory. Implicit parameters is another sigma parameter used both for 
% tubeness and OrientationJ
% Based on: Ofra Golani and G.Molodij 16/10/2017
% Modifyed for entropy measurement by Elee Shimshoni 12/11/2017
% Modifyed for calculation of mean entropy per mouse and then comparing by 
% Elee Shimshoni 15/1/2018.


clear all;
close all;

ymax=4; % set the value of ymax 

% After Fiji orientation analysis *.csv files for 91 values of orientations
% from -90 upto 90 degrees
ext='.csv';
numdata=91;
rep=1;
n=1;
maxfiles=500; %Maximum number of files of a sequence

% Start the plot that superpose selected sequences with different number of
% images

figure(1); hold on;
xplot=(1:10);
lim_orient(1:10)=1;
plot(xplot,lim_orient,'k-','LineWidth',2);
set(gca,'xticklabel',[]);
set(gca,'xtick',[]);


% Selection of the control sequences
prompt = 'How many control samples(type in number)?';
numCon = input(prompt);
for l=1:numCon
    sequences={};
    sample = uigetdir(pwd,'Select sample folder');
    sequences = struct2cell(dir(fullfile(sample, '*.tif')));
    s=strsplit(sample,'\');
    SampleNameCon{l}=s(end);
    numOfSeqCon(l)=size(sequences,2);
    PathNameCon{l}=sequences(2,1);
    for n=1:size(sequences,2)
        FileNameCon{l,n}=sequences(1,n);
        resFolder='Results/';
        WorkPathCon{l,n} = strcat(PathNameCon{l},'\',resFolder);
        FileCon{l,n}=strcat(PathNameCon{l},'\',FileNameCon{l,n});
        Ima=strcat(char([FileCon{l,n}]));
        Iinfo=imfinfo(Ima);
        numofileCon{l,n}=length(Iinfo);
    end

end
% End selection of control sequences

% Start processing and plot the selected control sequences
    l=1;
for l=1:numCon
    entropyCon=[];
for k=1:numOfSeqCon(l) 
    nnumofileCon=[numofileCon{l,k}];
    nWorkPathCon=char(WorkPathCon{l,k});
    nFileNameCon=char(FileNameCon{l,k});

    for j=1:nnumofileCon
        num=num2str(j);
        ftoread = strcat(nWorkPathCon,nFileNameCon,num,ext);
        fid = fopen(ftoread);
        sizex=[91];
        sizey=[91];
        for i=1:90
            fgetl(fid);
            M = textscan(fid, '%f, %f ,%f, %f,\n ');
            M=M';
            M=cell2mat(M);
            % After Fiji orientation analysis, M(3) is the raw value, M(4) 
            freqCon(i)=M(3);
        end
        fclose(fid)
        entropyCon(k,j)=-sum(freqCon.*log(freqCon));
    
    end
         entropyCon(entropyCon==0)=NaN;
         meanSeqEntropyCon(l,k)=nanmean(entropyCon(k,:)); %collects means for all images of a sample
    
end
        meanSeqEntropyCon(meanSeqEntropyCon==0)=NaN;
        meanEntropyCon(l)=nanmean(meanSeqEntropyCon(l,:)); %creates a vector of means of each sample
end
  
    allMeanEntropyCon=mean(meanEntropyCon);
    xplot=2;  % plot all control results at x = 2
    plot(xplot,allMeanEntropyCon,'k+','MarkerSize',20,'LineWidth',2);
    plot(xplot,meanEntropyCon,'b.','MarkerSize',10);

% Selection of the treated sequences
prompt = 'How many treated samples(type in number)?';
numTre = input(prompt);
for l=1:numTre
    sequences={};
    sample = uigetdir(pwd,'Select sample folder');
    sequences = struct2cell(dir(fullfile(sample, '*.tif')));
    s=strsplit(sample,'\');
    SampleNameTre{l}=s(end);
    numOfSeqTre(l)=size(sequences,2);
    PathNameTre{l}=sequences(2,1);
    for n=1:size(sequences,2)
        FileNameTre{l,n}=sequences(1,n);
        resFolder='Results\';
        WorkPathTre{l,n} = strcat(PathNameTre{l},'\',resFolder);
        FileTre{l,n}=strcat(PathNameTre{l},'\',FileNameTre{l,n});
        Ima=strcat(char([FileTre{l,n}]));
        Iinfo=imfinfo(Ima);
        numofileTre{l,n}=length(Iinfo);
    end

end
% End selection of treated sequences

% Start processing and plot the selected treated sequences
    l=1;
for l=1:numTre
    entropyTre=[];
for k=1:numOfSeqTre(l) 
    nnumofileTre=[numofileTre{l,k}];
    nWorkPathTre=char(WorkPathTre{l,k});
    nFileNameTre=char(FileNameTre{l,k});

    for j=1:nnumofileTre
        num=num2str(j);
        ftoread = strcat(nWorkPathTre,nFileNameTre,num,ext);
        fid = fopen(ftoread);
        sizex=[91];
        sizey=[91];
        for i=1:90
            fgetl(fid);
            M = textscan(fid, '%f, %f ,%f, %f,\n ');
            M=M';
            M=cell2mat(M);
            % After Fiji orientation analysis, M(3) is the raw value, M(4) 
            freqTre(i)=M(3);
        end
        fclose(fid)
        entropyTre(k,j)=-sum(freqTre.*log(freqTre));
    
    end
         entropyTre(entropyTre==0)=NaN;
         meanSeqEntropyTre(l,k)=nanmean(entropyTre(k,:)); %collects means for all images of a sample
    
end
        meanSeqEntropyTre(meanSeqEntropyTre==0)=NaN;
        meanEntropyTre(l)=nanmean(meanSeqEntropyTre(l,:)); %creates a vector of means of each sample
end
    
    allMeanEntropyTre=mean(meanEntropyTre);
    xplot=3;  % plot all treated results at x = 3
    plot(xplot,allMeanEntropyTre,'k+','MarkerSize',20,'LineWidth',2);
    plot(xplot,meanEntropyTre,'r.','MarkerSize',10); 

% Finish the plot that superpose selected sequences with different number of
% images

E=[meanEntropyCon,meanEntropyTre];
ymax=max(E)+1;
ymin=min(E)-1;
ylim([ymin ymax]);
xlim([1 4]);
ylabel('Entropy of distribution')
txt=texlabel('Control')
text(1.75,ymin-0.2,txt,'fontsize',18,'color','blu');
txt=texlabel('Treated')
text(2.75,ymin-0.2,txt,'fontsize',18,'color','red');
[h,p]=ttest2(meanEntropyCon,meanEntropyTre)

grid on;
hold off;

prompt = 'Do you want to remove outliers 1/0 (Yes/No)? ';
    rep = input(prompt);
    if rep == 1
    %create fig 2
    figure(2); hold on;
    xplot=(1:10);
    lim_orient(1:10)=1;
    plot(xplot,lim_orient,'k-','LineWidth',2);
    set(gca,'xticklabel',[]);
    set(gca,'xtick',[]);
  %calculate MAD  
      medCon=repmat((nanmedian(meanSeqEntropyCon'))',1,size(meanSeqEntropyCon,2));
      medTre=repmat((nanmedian(meanSeqEntropyTre'))',1,size(meanSeqEntropyTre,2));
      forMadCon=abs(meanSeqEntropyCon-medCon);
      forMadTre=abs(meanSeqEntropyTre-medTre);
      MADCon=repmat(1.4826*(nanmedian(forMadCon'))',1,size(meanSeqEntropyCon,2));
      MADTre=repmat(1.4826*(nanmedian(forMadTre'))',1,size(meanSeqEntropyTre,2));
  %identify outliers
  prompt = 'What cutoff factor do you want for outlier detection 2/3? ';
    rep = input(prompt);
        cutOff=rep;
        UpperCon=medCon+(cutOff*MADCon);
        LowerCon=medCon-(cutOff*MADCon);
        UpperTre=medTre+(cutOff*MADTre);
        LowerTre=medTre-(cutOff*MADTre);
        OutlierCon=or(meanSeqEntropyCon<LowerCon,meanSeqEntropyCon>UpperCon);
        OutlierTre=or(meanSeqEntropyTre<LowerTre,meanSeqEntropyTre>UpperTre);
        meanSeqEntropyConNO=meanSeqEntropyCon;
        meanSeqEntropyConNO((OutlierCon==1))=NaN;
        meanSeqEntropyTreNO=meanSeqEntropyTre;
        meanSeqEntropyTreNO((OutlierTre==1))=NaN;
        meanEntropyConNO=nanmean(meanSeqEntropyConNO');
        meanEntropyTreNO=nanmean(meanSeqEntropyTreNO');
        allMeanEntropyConNO=mean(meanEntropyConNO);
        allMeanEntropyTreNO=mean(meanEntropyTreNO);

  %plot
        xplotC=2;
        xplotT=3;
        plot(xplotC,allMeanEntropyConNO,'k+','MarkerSize',20,'LineWidth',2);
        plot(xplotC,meanEntropyConNO,'b.','MarkerSize',10);
        plot(xplotT,allMeanEntropyTreNO,'k+','MarkerSize',20,'LineWidth',2);
        plot(xplotT,meanEntropyTreNO,'r.','MarkerSize',10); 
        E=[meanEntropyConNO,meanEntropyTreNO];
        ymax=max(E)+1;
        ymin=min(E)-1;
        ylim([ymin ymax]);
        xlim([1 4]);
        ylabel('Entropy of distribution')
        txt=texlabel('Control')
        text(1.75,ymin-0.2,txt,'fontsize',18,'color','blu');
        txt=texlabel('Treated')
        text(2.75,ymin-0.2,txt,'fontsize',18,'color','red');
%t test
        [h,pNO]=ttest2(meanEntropyConNO,meanEntropyTreNO)


    hold off;

    prompt = 'Do you want to change cutoff factor? 1/0 (Yes/No)? ';
    rep = input(prompt);
    if rep == 1
         prompt = 'What cutoff factor do you want for outlier detection 2/3? ';
    rep = input(prompt);
    %create fig 3
    figure(3); hold on;
    xplot=(1:10);
    lim_orient(1:10)=1;
    plot(xplot,lim_orient,'k-','LineWidth',2);
    set(gca,'xticklabel',[]);
    set(gca,'xtick',[]);
    %detect outliers    
        cutOff=rep;
        UpperCon=medCon+(cutOff*MADCon);
        LowerCon=medCon-(cutOff*MADCon);
        UpperTre=medTre+(cutOff*MADTre);
        LowerTre=medTre-(cutOff*MADTre);
        OutlierCon=or(meanSeqEntropyCon<LowerCon,meanSeqEntropyCon>UpperCon);
        OutlierTre=or(meanSeqEntropyTre<LowerTre,meanSeqEntropyTre>UpperTre);
        meanSeqEntropyConNO=meanSeqEntropyCon;
        meanSeqEntropyConNO((OutlierCon==1))=NaN;
        meanSeqEntropyTreNO=meanSeqEntropyTre;
        meanSeqEntropyTreNO((OutlierTre==1))=NaN;
        meanEntropyConNO=nanmean(meanSeqEntropyConNO');
        meanEntropyTreNO=nanmean(meanSeqEntropyTreNO');
        allMeanEntropyConNO=mean(meanEntropyConNO);
        allMeanEntropyTreNO=mean(meanEntropyTreNO);
   
  %plot
        xplotC=2;
        xplotT=3;
        plot(xplotC,allMeanEntropyConNO,'k+','MarkerSize',20,'LineWidth',2);
        plot(xplotC,meanEntropyConNO,'b.','MarkerSize',10);
        plot(xplotT,allMeanEntropyTreNO,'k+','MarkerSize',20,'LineWidth',2);
        plot(xplotT,meanEntropyTreNO,'r.','MarkerSize',10); 
        E=[meanEntropyConNO,meanEntropyTreNO];
        ymax=max(E)+1;
        ymin=min(E)-1;
        ylim([ymin ymax]);
        xlim([1 4]);
        ylabel('Entropy of distribution')
        txt=texlabel('Control')
        text(1.75,ymin-0.2,txt,'fontsize',18,'color','blu');
        txt=texlabel('Treated')
        text(2.75,ymin-0.2,txt,'fontsize',18,'color','red');
%t test
        [h,pNO]=ttest2(meanEntropyConNO,meanEntropyTreNO)

    hold off;
 
    end
    end
    
```
