## supplemental text and figures S1-S6 for "Enhancing Uterine Receptivity for Embryo Implantation through Controlled Collagenase Intervention"

**This PDF file includes:**
Supplementary Text
Figs. S1 to S7

**Other Supplementary Materials for this manuscript include the following:**
Two macrocode files named “Orientation” and “Entropy”

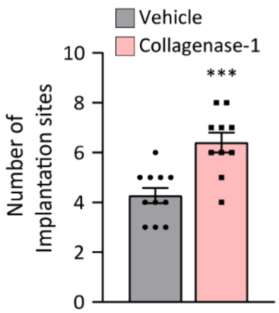


**Fig. S1. Treatment with collagenase-1 improves implantation rate *in-vivo*.** Number of implantation sites in the spontaneous pregnancy model using the C57BL/6 mouse strain (n≥10). Data were analyzed by an unpaired, two-tailed t-test. Results are presented as mean ± SEM with significance: ***p < 0.001.

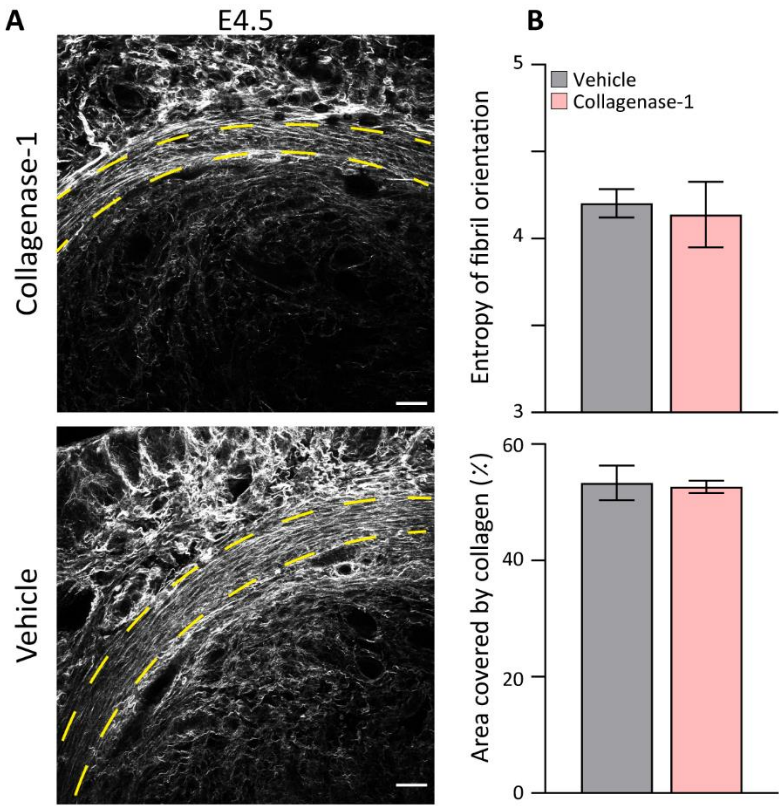


**Fig. S2. Treatment with collagenase-1 does not affect organization of the myometrium collagen fibers. (A)** Representative SHG images of cross sections of vehicle- and collagenase-1 treated uterine samples at E4.5. Yellow dashed lines represent the myometrium layer (scale bar= 50 µM). **(B)** Measurements of the entropy of fiber orientation (upper), and area covered by collagen (lower). Analysis was done using ImageJ software. Resulting orientation data were analyzed using Matlab code to assess fiber entropy (n=2).

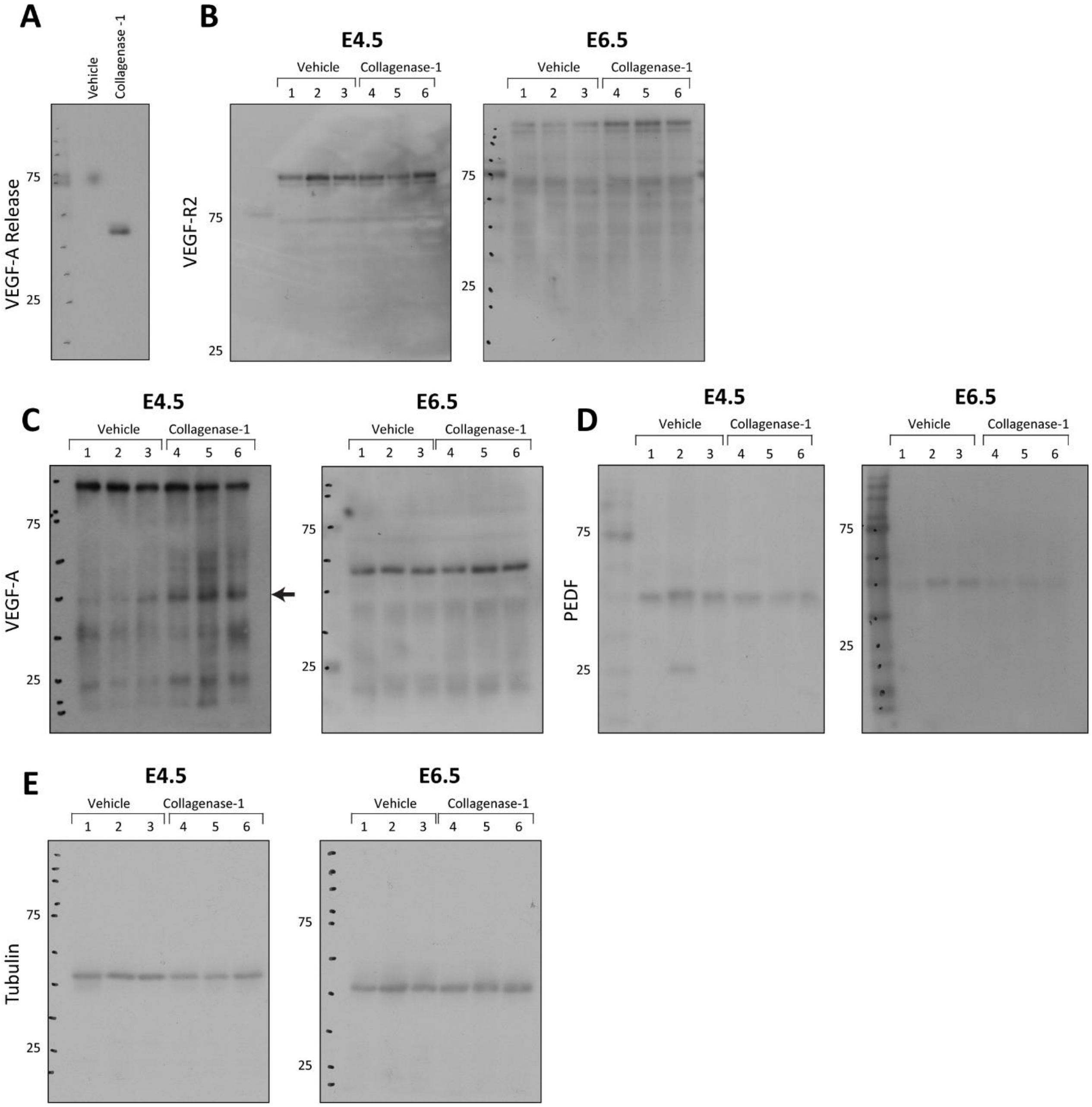


**Fig. S3. The full membranes of western blots presented in Figures 4 and 5**, showing expression of the indicated proteins.

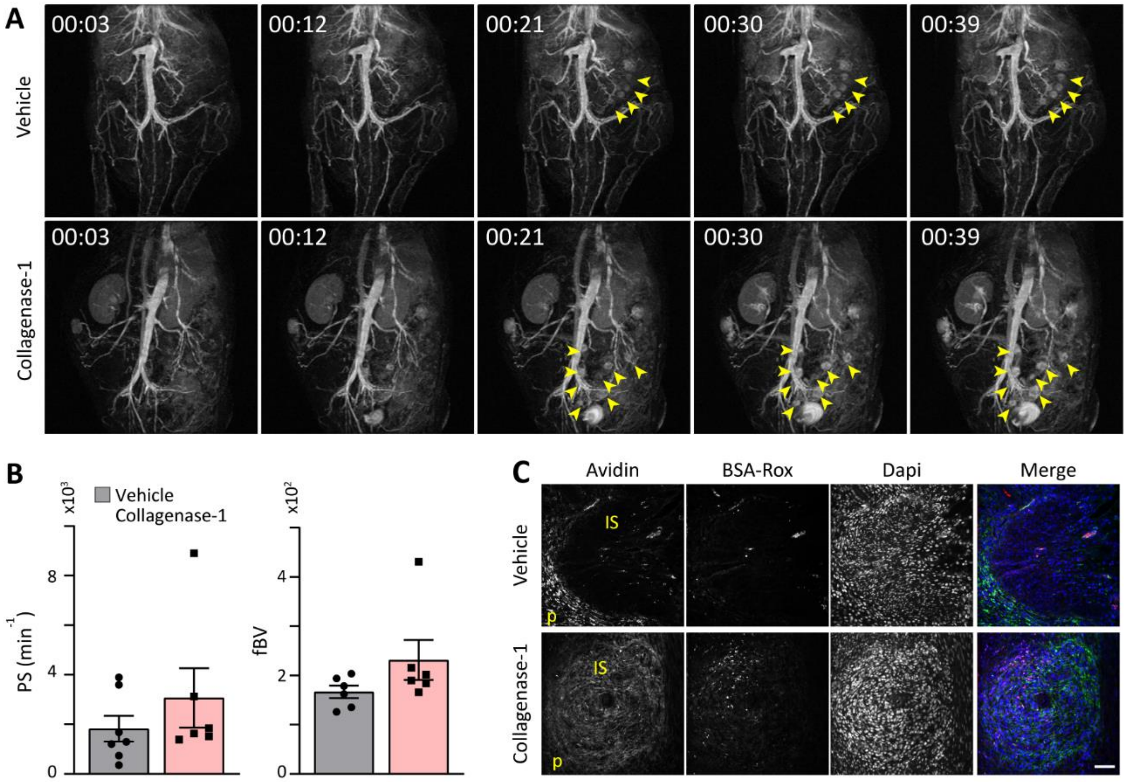


**Fig. S4. Collagenase-1 promotes vascular permeability at implantation sites**

**(A)** Representative MRI images at E4.5 of vehicle- and collagenase-1-treated uteri. Arrows indicate implantation sites. **(B)** Quantification of the permeability surface (PS) and fractional blood volume (fbv) at implantation sites, taken from the MRI data (n = 7; 9.4T, 3D GE; IV biotin-BSA-GdDTPA). **(C)** Immunofluorescence imaging of the contrast agents: biotin-BSA-Gd-DTPA (green) and BSA-Rox (red), demonstrating their accumulation in the implantation sites during MRI imaging for 30 and 2.5 min, respectively.

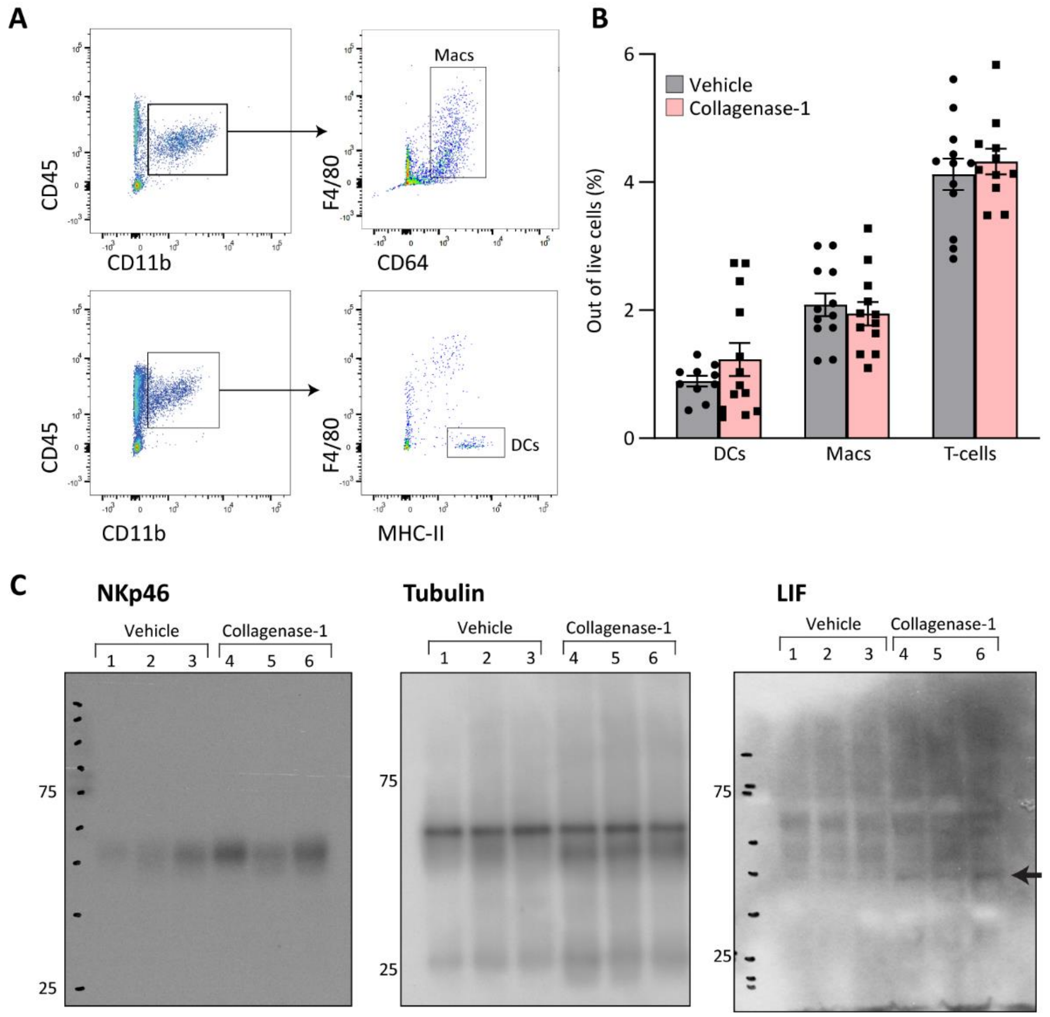


**Fig. S5. Collagenase-1 induces upregulation of the cytokine LIF and NK cell infiltration in the spontaneous pregnancy model. (A)** Representative flow cytometry images showing the gating strategy used to identify macrophages (CD45+CD11b+F4/80+CD64+) and dendritic cells (CD45+CD11c+F4/80−MHC-II+). Macs, macrophages (upper panel). DCs, dendritic cells (lower panel). **(B)** Quantification of flow cytometry results showing the percent of macrophages, dendritic cells, and T cells among live cells at E4.5 (n ≥ 12). Data were analyzed by an unpaired, two-tailed t-test. Results are presented as mean ± SEM. **(C)** Western blot full membranes of indicated proteins presented in Fig. 6C. Arrow indicates the specific bend for LIF.

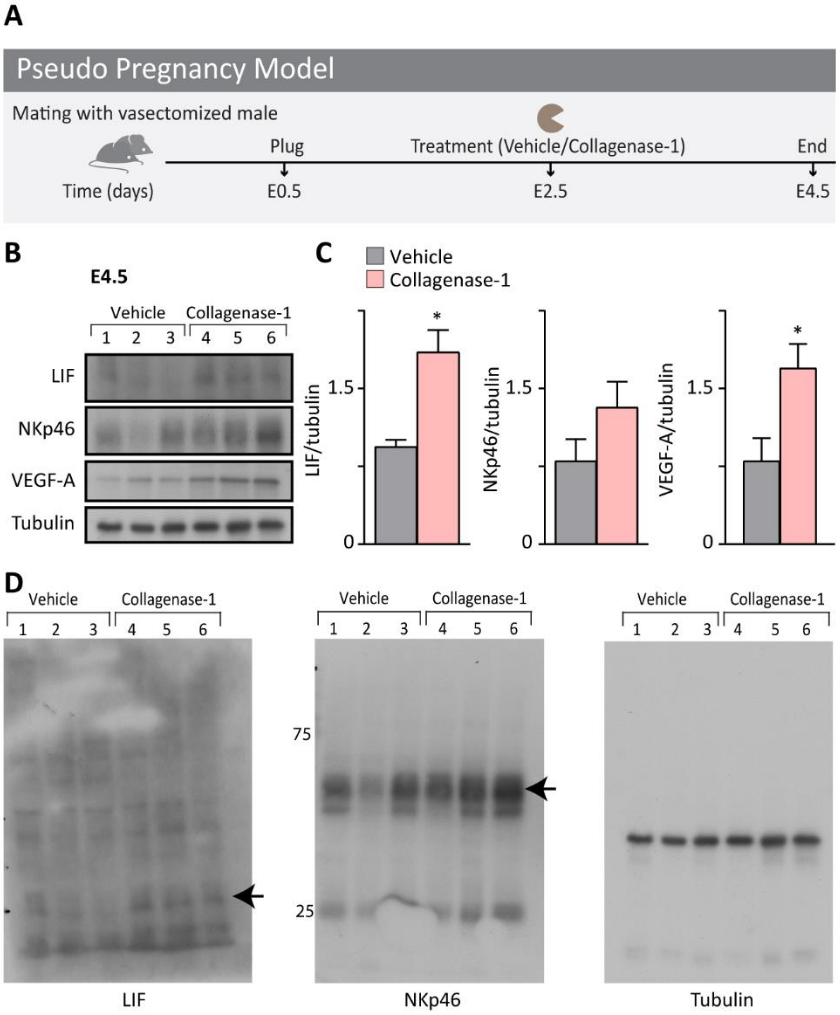


**Fig. S6. Collagenase-1 induces inflammatory and angiogenesis-related pathways independent of embryos presence.** **(A)** Schematic representation of the pseudo pregnancy experimental model. **(B)** Western blot analysis of E2.5 samples treated with either collagenase-1 or vehicle. **(C)** Quantification of western blots using ImageJ analysis tool (n=3). **(D)** Full membranes of indicated proteins presented in Figure 6. Arrow indicates the protein's specific bend. Data were analyzed by an unpaired, two-tailed t-test. Results are presented as mean ± SEM with significance: *$p<0.05$, **$p < 0.01$.

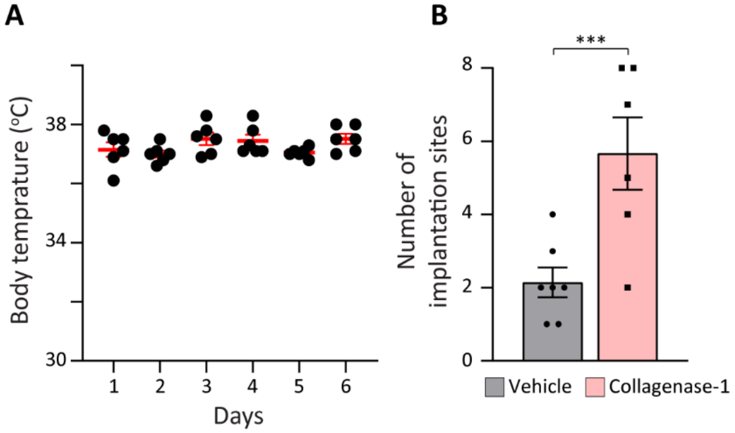


**Fig. S7. Body temperature during the heat-stress protocol and improvement of implantation rate in a cross-strain embryo transfer model. (A)** Monitoring of mice body temperature during the heat-stress protocol, ranged between 36 ºC – 38 ºC throughout the experiment, indicating that females were in a normal and healthy physiological state. **(B)** Number of implantation sites in the cross-strain embryo transfer model in which cbcF1 embryos were transferred to pseudo-pregnant ICR females with or without collagenase-1 treatment ($n \geq 6$). Data were analyzed by an unpaired, two-tailed t-test. Results are presented as mean ± SEM with significance: **$p < 0.01$.
